# Single-point mutation alters odorant receptor sensitivity associated with host plant specialization in *Spodoptera* moths

**DOI:** 10.64898/2026.09.24.754037

**Authors:** Arthur Comte, Maxence Lalis, Sai Zhang, Albert Thorhallsson, Riccardo Moracci, Jérémy Gévar, Nicolas Montagné, Jérémie Topin, Andrew Mitchell, Sébastien Fiorucci, Emmanuelle Jacquin-Joly

## Abstract

Host specialization in herbivorous insects is often associated with divergence in chemosensory abilities. Here, we investigated the possible contribution of odorant receptors (ORs) in host plant restriction in the lily moth *Spodoptera picta*, a species specialized on Amaryllidaceae. Manual annotation of *S. picta* ORs in its genome revealed a repertoire similar in size and composition to those of its polyphagous sister species, *S. littoralis* and S. *litura*, suggesting that specialization did not involve major gene loss or expansion in the lily moth. To assess functional divergence beyond gene number, we applied a large scaled structure-based virtual screening approach to the entire OR repertoires of these three *Spodoptera* species, generating ligand-binding profiles for 120,591 volatile compounds. Among 69 1:1:1 OR orthologs, 24 exhibited divergent predicted binding spectra. We pinpointed OR29 that we also found to be highly expressed in both male and female antennae of *S. picta* through a RNAseq approach. Functional assays demonstrated that *S. picta* OR29 acquired heightened sensitivity to limonene enantiomers, volatiles emitted by host Amaryllidaceae inflorescences. Site-directed mutagenesis revealed that a single amino acid substitution within the predicted binding region underlies this shift in sensitivity. These results show that host specialization in *S. picta* has not been accompanied by significant OR repertoire remodeling, but rather by subtle molecular changes that fine-tune receptor sensitivity to host-derived volatiles.

## Introduction

Host specialization in insects is accompanied by physiological (Zdenek et al. 2024), morphological (Ceja-Navarro et al. 2019), and behavioral adaptations (Álvarez-Ocaña et al. 2023) that facilitate the exploitation of novel food sources. Among these, changes in the digestive system are well documented, including structural modifications of the mouthparts (Bauder and Karolyi 2019) that enhance nutrient extraction, and the evolution of specialized detoxification enzymes (Gikonyo et al. 2024) which enables the neutralization of host defensive compounds. However, successful colonization of a host also depends on the insect’s ability to detect, identify, and navigate toward appropriate targets, tasks that are partly governed by the olfactory system (Montagné et al. 2022). Olfactory adaptations therefore play a role in initiating or stabilizing host specialization by reshaping the insect’s responsiveness to host-specific chemical cues (Andersson et al. 2015).

In neopteran insects, the olfactory process begins with the peripheral detection of odorant compounds, which is primarily mediated by a family of seven-transmembrane domain (7-TM) proteins (Clyne et al. 1999; Gao and Chess 1999; Vosshall et al. 1999) known as odorant receptors (ORs). These receptors are expressed in the dendritic membrane of olfactory sensory neurons (OSNs) housed within specialized olfactory organs, mainly the antennae. ORs form heterotetrameric ion channel complexes (Wang et al. 2024b; Zhao et al. 2024) with an obligatory co-receptor, Orco (Larsson et al. 2004). The OR-Orco complexes translate chemical cues from the environment into electrical signals, which are then relayed to the central nervous system for information processing. The OR subunits of the complex, which contain the odorant-binding site, are highly divergent both between and within species, and impart chemical specificity to the complex. In contrast, Orco is highly conserved in neopteran insects and essential for proper subcellular trafficking of ORs (Larsson et al. 2004), as well as for shaping the architecture of the ion channel pore, alongside the variable OR subunits (Wang et al. 2024b; Zhao et al. 2024). Through high-resolution cryo-electron microscopy (cryo-EM), a series of apo and ligand-bound structures of neopteran insect olfactory ion channels has been recently resolved, comprising six distinct OR–Orco complexes from representative species of Hemiptera (Wang et al. 2024b; Dong et al. 2026), Diptera (Zhao et al. 2024; Wang et al. 2026), and Lepidoptera (Jang et al., unpublished data).

The OR gene family is characterized by rapid evolutionary dynamics, shaped by processes of gene duplication, sequence divergence, pseudogenization, and loss—a pattern known as “birth-and-death” evolution (Robertson 2019). These dynamics can lead to lineage-specific expansions or contractions of OR repertoires, often aligning with major ecological transitions. For instance, in the herbivorous Drosophilidae *Scaptomyza flava* that evolved from microbe-feeding ancestors, the shift to feeding and ovipositing on living plant tissue coincided with the loss of yeast-volatile-detecting ORs and the duplication of an OR tuned to green leaf volatiles (Goldman-Huertas et al. 2015). At broader phylogenetic scales, comparative studies reported positive correlation between host breadth and OR repertoire size (Robertson et al. 2018; Andersson et al. 2019). However, the correlation weakens when comparisons are made within smaller phylogenetic scales, such as families (He et al. 2018). Comparative studies of closely related species suggested that, although gene family expansion and contraction can reshape the chemical space detected by organisms and contribute to host specialization, other mechanisms affecting the OR repertoire—such as differential gene expression or functional divergence among orthologs—may play a more prominent role. For instance, the host specialization of the domestic form of the mosquito *Aedes aegypti* from non-human animals to humans along the Kenyan coast has been associated with increased expression and heightened ligand sensitivity of the odorant receptor OR4, which is specifically tuned to sulcatone—a volatile compound enriched in human odor (McBride et al. 2014). As another example, a single-point mutation within the binding pocket of OR22a has altered the receptor’s tuning toward methyl esters during the evolution of *Drosophila sechellia*, contributing to this species’ specialization on noni fruit compared to its sister species, *D. melanogaster* (Auer et al. 2020). These studies, focused on species from the Diptera order, provide key insights into the complex mechanisms driving OR adaptation during host specializations. However, broadening comparative analyses to include other insect orders and fine analyses of the molecular bases of such transitions are essential to fully understand the relationship between OR evolution and host selection.

Within Lepidoptera, the genus *Spodoptera* comprises 31 species of noctuid moths that exhibit diverse host plant ranges (Kergoat et al. 2021). While the chemical ecology of several *Spodoptera* species, such as *S. litura*, *S. littoralis,* and *S. frugiperda*, have been extensively studied due to their economic importance as agricultural pests, the ecology of others—such as *S. picta*—has been neglected. In contrast to its two sister species *S. litura* and *S. littoralis*, *S. picta* exhibits an unusually narrow host range (Fig. 1), with larvae found exclusively on certain plants within the Amaryllidaceae family (Ang et al. 2010). Phylogenetic analyses suggest that *S. picta*, *S. litura,* and *S. littoralis* diverged from a common ancestor approximately 4.45 million years ago (Kergoat et al. 2021) (Fig. 1). A recent study investigated its host specialization by analyzing its detoxification systems, revealing a loss of key detoxification enzymes compared to other *Spodoptera* species (Wang et al. 2024a). However, whether olfactory divergence is linked to host specialization in *S. picta* compared to its sister species remains unexplored.

**Fig. 1.**
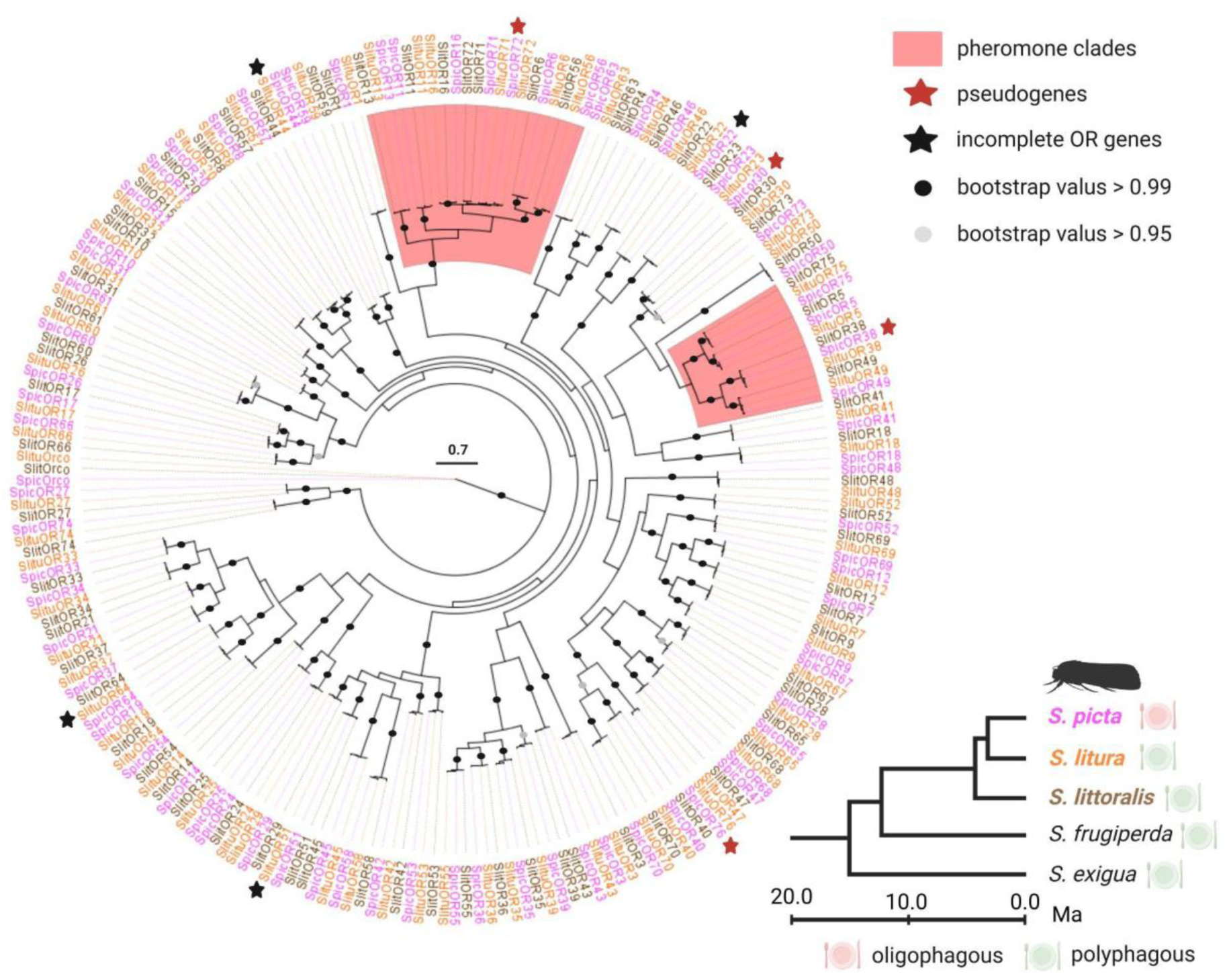
The OR repertoire of *Spodoptera picta* is similar to those of its sister species with broader host ranges, *S. litura* and *S. littoralis*. Phylogenetic tree of *S. picta* (Spic), *S. litura* (Slitu), and *S. littoralis* (Slit) ORs, rooted with Orco sequences. SpicORs are represented in pink, SlituORs in orange, and SlitORs in brown. Among the three species, 99% of ORs show a clear 1:1:1 orthology relationship. Pseudogenes are marked with red stars, whereas incomplete genes are indicated by black stars. Clusters that include known *Spodoptera* sex pheromone receptors are highlighted in red. Only bootstrap values above 95 are shown: black dots indicate values greater than 99, while grey dots indicate values between 95 and 99. The scale bar represents 0.7 amino acid substitutions per site. Sequences were aligned using MAFFT 7 web server (Katoh et al. 2019). The phylogenetic tree was reconstructed using the PhyML 3.0 webserver (Guindon et al. 2005) and visualized with FigTree 1.4.4 (https://github.com/rambaut/figtree/releases/tag/v1.4.4). A truncated phylogenetic tree of the genus *Spodoptera*, including the studied species, is shown in the bottom-right corner of the plot. Evolutionary relationships and divergence times are based on Kergoat et al. (2021). Labels indicate dietary breadth (oligophagous or polyphagous) for each species.

In this study, we first annotated the complete OR repertoire in the available *S. picta* genome (Wang et al. 2024a). Next, we combined a predictive definition of each OR chemical space with transcriptomic data to identify candidate *S. picta* ORs (further referred as SpicORs) potentially associated with host specialization. This integrative approach led us to identify SpicOR29, a receptor highly expressed in the antennae of *S. picta*, which was found to exhibit distinct sensitivity to volatiles emitted by the Amaryllidaceae host plants (*Clivia miniata* and *Crinum asiaticum*) compared to its orthologs in species with broader host ranges. Through site-directed mutagenesis, we showed that a single-point mutation targeting a residue within the predicted binding region underlies this functional divergence.

## Results

### Annotation of *S. picta* odorant receptors: no major contraction nor expansion in OR gene repertoire compared to sister species with broader host ranges

We first determined the full repertoire of SpicORs in the published *S. picta* genome (GenBank accession GCA_038387885.1; Wang et al. 2024a) using alignment of OR sequences from *S. exigua*, *S. frugiperda*, *S. litura*, and *S. littoralis*, followed by manual curation of gene models. We annotated a total of 75 SpicOR genes, including the conserved co-receptor Orco (supplementary table S1). Four genes (SpicOR30, SpicOR38, SpicOR72, and SpicOR76) were classified as pseudogenes based on the presence of premature stop codons or frameshift-inducing insertions and deletions that disrupted the open reading frame; while one additional gene, SpicOR22, was partially annotated due to limitations in the genome assembly. The remaining 69 SpicOR genes were full-length and intact. The SpicOR repertoire resembled those of its sister species, *S. litura* and *S. littoralis*, which contain 75 and 74 OR genes, respectively, including Orcos. No major contraction or expansion was observed, with over 98% of ORs showing a 1:1:1 orthology relationship among the three species (Fig. 1). We identified nine intact *S. picta* ORs that clustered in known *Spodoptera* sex pheromone receptor clades, representing candidate pheromone receptors in this species: SpicOR5, 6, 11, 13, 16, 49, 56, 71, and 75 (Fig. 1).

### Prediction of functionally divergent orthologous ORs across *Spodoptera* species

To identify orthologous ORs capable of detecting distinct chemicals between *S. picta* and its sister species, we computed a functional divergence distance based on an OR-molecule interaction fingerprint established for each OR. These fingerprints represent predicted interactions between each receptor and a set of 120,591 natural volatile products, determined through molecular docking simulations using a previously validated protocol established on *S. littoralis* ORs (Comte et al. 2025). Three-dimensional structures of 69 SpicORs, 71 SlituORs, and 73 SlitORs were predicted with AlphaFold2 (Jumper et al. 2021), excluding Orcos, pseudogenes, and incomplete sequences. Dimensionality of each interaction fingerprint was subsequently reduced using principal component analysis (PCA), followed by a HDBSCAN clustering (Campello et al. 2013) based on Euclidean distances to classify the receptors according to their predicted chemical detection profiles. Among the 69 SpicORs modeled, 24 did not cluster with their respective orthologs from *S. litura* and *S. littoralis* (namely, SpicOR3, 4, 6, 7, 14, 18, 19, 23, 25, 26, 28, 29, 31, 34, 45, 52, 53, 56, 57, 65, 67, 68, 69 and 74). Ranking these receptors according to the minimal distance to their orthologs revealed that SpicOR23, SpicOR65, SpicOR29, SpicOR68 and SpicOR28 exhibited the greatest predicted functional divergence (Fig. 2a and c).

**Fig. 2.**
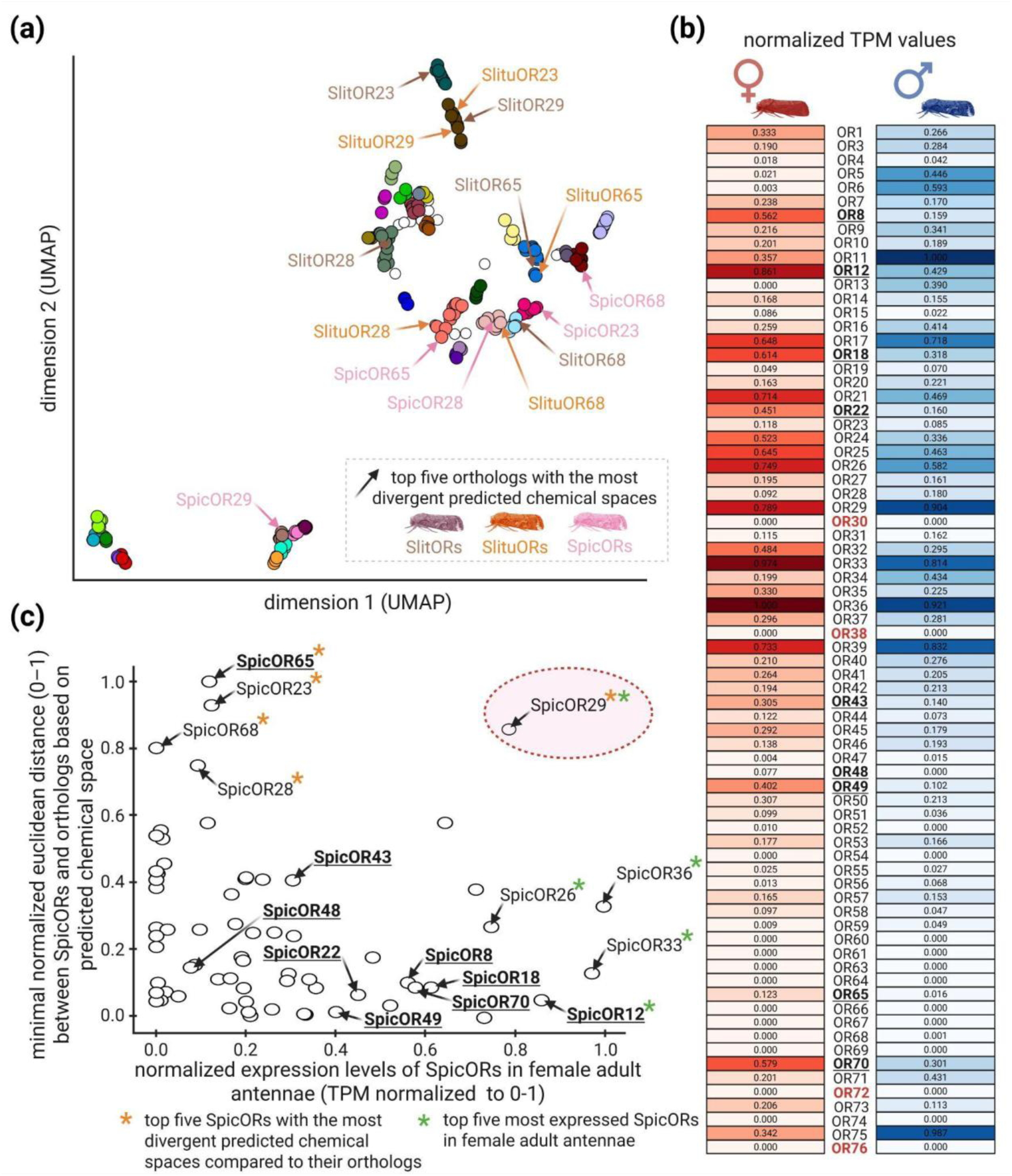
SpicOR29 emerges as a promising candidate for olfactory divergence between *S. picta*, *S. litura*, and *S. littoralis*. Its predicted chemical space is highly divergent from that of its orthologs, and it is strongly expressed in the antennae of virgin adult females. (**a**) Uniform manifold approximation and projection (UMAP) (McInnes et al. 2018) representation of SpicOR, SlituOR, and SlitOR predicted chemical spaces based on the reevaluated Vinardo docking scores (Comte et al. 2025) for 120,591 docked molecules. Each colored dot represents a clustered OR, with distinct clusters identified by HDBSCAN (Campello et al. 2013) and represented by different colors. Unclustered ORs are shown as white dots. The top five of the SpicORs with the most divergent predicted chemical spaces compared to their orthologs are highlighted in the graph. The corresponding SpicORs and their orthologs are indicated by arrows, with protein names provided and color-coded by species (SpicORs: pink, SlituORs: orange; SlitORs: brown). (**b**) Heatmap displaying the mean TPM values of SpicOR expression, normalized between 0 and 1, across three replicates in the antennae of virgin adult females (red) and virgin adult males (blue). The intensity of the color corresponds to expression levels, with darker shades indicating higher TPM values. Gene names showing a significant difference in expression between sexes (DESeq2 with Benjamini-Hochberg correction; padj < 0.1) are bold and underlined. Pseudogenes are bold and colored in red. (**c**) Plot showing the relationship between the minimal normalized euclidean distance between SpicORs and their orthologs, based on their predicted chemical space (Y-axis), and the expression levels of SpicORs in virgin adult female antennae (TPM) (X-axis). Both values are normalized to a scale of 0–1. Orange asterisks highlight the top five SpicORs exhibiting the highest divergence in predicted chemical space compared to their orthologs. Green asterisks indicate the SpicORs with the highest expression levels in virgin adult female antennae. Gene names showing a significant difference in expression between sexes (DESeq2 with Benjamini-Hochberg correction; padj < 0.1) are bold and underlined. The position of SpicOR29 is highlighted in red.

### SpicOR29: an OR combining high expression in *S. picta* antennae and a predicted divergent chemical space compared to its *S. litura* and *S. littoralis* orthologs

To refine the list of ORs possibly involved in host choice, we combined the functional divergence distance with gene expression data. We hypothesized that the ORs associated with host specialization in *S. picta* would not only display distinct predicted chemical spaces compared to their orthologs, but also elevated expression levels in the antennae of adult females. Indeed, previous functional studies in moths have demonstrated that female-biased (Liu et al. 2020; Wang et al. 2023b) or highly antennally expressed ORs (Ma et al. 2024) contribute to host selection.

To measure the expression levels of SpicORs, we performed RNA-seq analysis. Illumina sequencing was performed on six libraries, three from *S. picta* virgin adult male antennae and three from virgin adult female antennae. Following filtering low-quality raw reads, each library contained between 50 and 56 million clean reads. The six datasets were assembled in a reference transcriptome. After redundant sequence clustering, the final reference transcriptome consisted of 45,622 unigenes. The BUSCO analysis showed a satisfying level of completeness and a low level of redundancy, with approximately 79% of BUSCO genes identified as complete and in a single-copy.

We identified 62 OR transcripts in addition to Orco in the reference antennal transcriptome (Table S3). Transcripts were not detected for SpicOR30, 38, 54, 60, 61, 63, 64, 66, 67, 69, 72, 74, and 76, including the four SpicORs predicted as pseudogenes in the genomic annotation. Notably, with the exception of the pseudogenes SpicOR38 and SpicOR72, none of these non-expressed SpicORs were assigned to the pheromone clades (Fig. 1). The number of ORs expressed in *S. picta* adult antennae was comparable to that reported for *S. litura* and *S. littoralis* in previous studies, which each express 60 ORs (Walker III et al. 2019; Yang et al. 2024). Of the 62 SpicOR transcripts, 39 encoded full-length ORs. Notably, the sequence of SpicOR22 was fully assembled in our transcriptome, in contrast to the truncated genomic sequence (supplementary table S1).

Nine ORs, SpicOR8, 12, 22, 43, 48, 49, 65, and 70, were significantly more expressed in adult female antennae than in adult male antennae (Fig. 2b). Among these, only SpicOR65 was predicted to be functionally divergent from its orthologs in *S. littoralis* and *S. litura*, but it was not retained for further analyses due to its low expression level. Based on Transcripts Per Million (TPM) values, SpicOR36, SpicOR33, SpicOR12, SpicOR29, and SpicOR26 were identified as the top five most highly expressed ORs in the antennae of *S. picta* adult females (Fig. 2b and c). SpicOR29 was the only receptor that ranked among both the five most highly expressed SpicORs in the antennae of *S. picta* adult females and the top five SpicORs with the most divergent predicted chemical spaces compared to their orthologs. Given this dual distinction, SpicOR29 appeared as the most promising OR for further studies. In the next steps, we thus focused on SpicOR29 and its orthologs, SlitOR29 and SlituOR29.

### OR29 orthologs present major functional differences according to sensitivity towards volatiles emitted by *S. picta* host plants

We next characterized and compared the recognition spectra of the three OR29 orthologs (SpicOR29, SlitOR29, and SlitOR29) using single-sensillum recordings on *Drosophila* ab3A OSNs expressing one of each SpicOR29, SlituOR29, and SlitOR29 in place of the endogenous DmelOR, an expression system known as the *Drosophila* “empty neuron system” (Chahda et al. 2019). The odorant panel was selected from floral volatiles emitted by *S. picta* Amaryllidaceae host plants, including *Clivia miniata* and *Crinum asiaticum* (Miyake et al. 1998; Kiepiel and Johnson 2014), retaining only compounds predicted by our previous structure-based virtual screening (SBVS) analysis to bind to at least one OR29 ortholog (Fig. 3a). Previously characterized SlitOR29 ligands were additionally included based on their predicted binding with SpicOR29 and SlituOR29 (de Fouchier et al. 2017) (Fig. 3a).

**Fig. 3.**
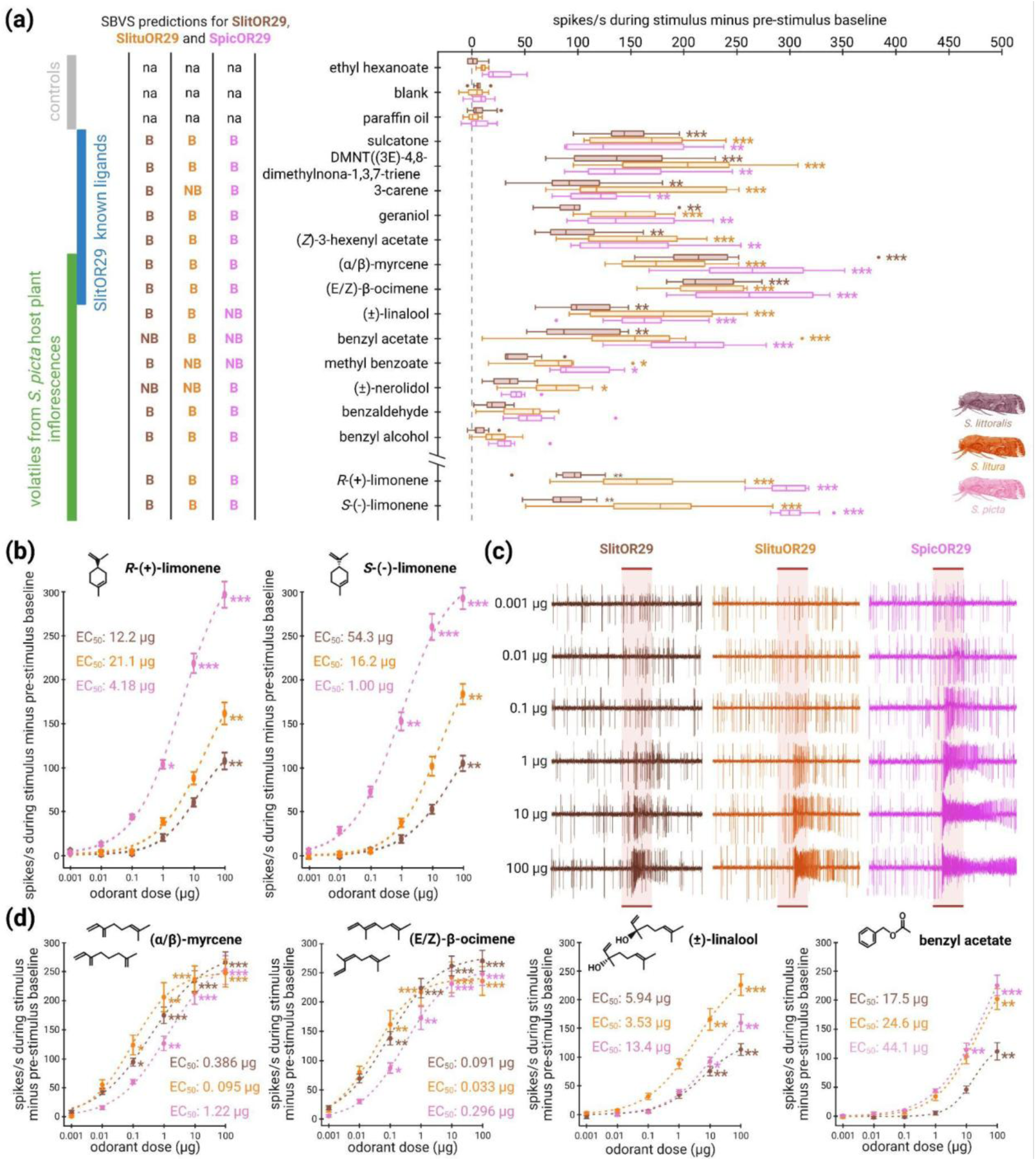
The *S. picta*, *S. litura*, and *S. littoralis* OR29 orthologs exhibit contrasted sensitivity to some *S. picta* host plant volatiles. (**a**) Box plot showing the responses of *Drosophila* ab3A OSNs (n = 8 biological replicates, each corresponding to a single OSN from a different individual fly) expressing SpicOR29 (pink), SlituOR29 (orange) or SlitOR29 (brown), measured upon exposure to controls, known ligands of SlitOR29, and volatiles present in the headspace of *S. picta* host plant and predicted by our SBVS to bind to at least one of the OR29 orthologs. All compounds were tested at 100 µg loaded onto filter paper in the stimulus cartridge. Responses to limonene enantiomers are markedly distinct, revealing strong functional divergence between the OR29 orthologs. The whiskers are extended to 1.5 times the interquartile range from the quartiles. Outliers are indicated by dots. To the left of the box plots, a summary table reports the SBVS predictions for each OR29 ortholog. Predicted binders are denoted “B” and non-binders “NB”, with the corresponding ORs color-coded by species (SpicOR29: pink; SlituOR29: orange; SlitOR29: brown). (**b**) Dose-response curves of *Drosophila* ab3A OSNs expressing one of each OR29 orthologs in response to the two limonene enantiomers. (**c**) Representative traces recorded from *Drosophila* OSNs expressing SpicOR29 (pink), SlituOR29 (orange), or SlitOR29 (brown) in response to increasing doses of *R*-(+)-limonene. The 500 ms stimulation window is indicated by a red bar. (**d**) Dose-response curves of *Drosophila* ab3A OSNs expressing one of each OR29 orthologs in response to selected ligands other than the limonene enantiomers that elicited strong responses in the panel screening. In (b and d), data represented are mean action potential frequencies ± SEM (n = 8 biological replicates, each corresponding to a single OSN from a different individual fly). Color coding reflects the species origin of the expressed receptor: pink, *S. picta*; brown, *S. littoralis*; orange, *S. litura*. Dose-response data were fitted with sigmoid functions using the curve_fit function from the SciPy 1.16.0 library. EC₅₀ values were determined from the fitted sigmoid dose–response curves and are indicated for each curve. In (**a, b and d**), asterisks indicate statistically significant differences between responses to the odorant and the solvent (Friedman test, followed by a Dunn’s test with a Benjamini & Yekutieli correction; \**padj* < 0.05, \*\**padj* < 0.01, \*\*\**padj* < 0.001).

The three OR29 orthologs were functional in *Drosophila* ab3A neurons. All were characterized as broadly tuned receptors, responding to 11, 13, and 12 out of the 15 compounds tested, for SpicOR29, SlitOR29, and SlitOR29, respectively (Fig. 3a). SBVS predictions showed good predictive performance across the tested compounds, with accuracies of 73%, 67%, and 60% for SlitOR29, SlituOR29, and SpicOR29, respectively (Fig. 3a). These results support the robustness of our predictive workflow and provide confidence in the predicted chemical spaces of SlitORs, SlituORs, and SpicORs, which successfully guided the selection of OR29 orthologs for functional analyses. Whereas some ligands have already been described for SlitOR29 (de Fouchier et al. 2017), we identified four new ligands for this OR, extending its response spectrum. We also report the first identification of ligands for SpicOR29 and SlituOR29. OR29 orthologs were activated by several volatiles present in the headspace of *S. picta* host plant inflorescences, including (α/β)-myrcene, (*E*/*Z*)-β-ocimene, (±)-linalool, benzyl acetate, and limonene enantiomers (*R*-(+)- and *S*-(−)-limonene) (Fig. 3a, supplementary fig. S1a). Among these, (*E*/*Z*)-β-ocimene and (α/β)-myrcene elicited the strongest responses from ab3A OSNs expressing SlituOR29 or SlitOR29, reaching median frequencies of 223 and 222 spikes/s for (*E*/*Z*)-β-ocimene, and 183 and 228 spikes/s for (α/β)-myrcene, respectively. In contrast, neurons expressing SpicOR29 were most strongly activated by limonene enantiomers, with *R*-(+)- and *S*-(−)-limonene eliciting median spike frequencies of 295 and 304 spikes/s, respectively (Fig. 3a). Moreover, differences in the selectivity of the OR29 orthologs were observed for one compound released by flowering *S. picta* host plants: methyl benzoate was detected by SpicOR29 and SlituOR29 but not by SlitOR29 (Fig. 3a).

To go further in the functional characterization of OR29 orthologs, we explored their sensitivity to the ligands that elicited the strongest responses in the previous screen, by conducting dose-response experiments (Fig. 3b to d). The most striking difference in sensitivity among the OR29 orthologs was observed with the limonene enantiomers (Fig. 3b and c). SpicOR29 was 100 times more sensitive to both *R*-(+)-limonene and *S*-(−)-limonene than SlitOR29 and SlituOR29, with significant responses starting at 1 µg on the filter paper, whereas activation of SlituOR29 and SlitOR29 required as much as 100 µg (Fig. 3b). The three OR29 orthologs exhibited similar sensitivities to (*E*/*Z*)-β-ocimene and (±)-linalool, activating *Drosophila* OSNs at doses starting from 0.1 µg and 10 µg, respectively (Fig. 3d). By contrast, SpicOR29 was 10 times less sensitive to (α/β)-myrcene than SlituOR29 and SlitOR29, with OSN activation starting at 1 µg compared to 0.1 µg for the other two receptors (Fig. 3d), and SlitOR29 was 10 times less sensitive to benzyl acetate than SpicOR29 and SlituOR29, requiring 100 µg to elicit a response, whereas the other orthologs responded at 10 µg (Fig. 3d).

### Single-point mutation alters the sensitivity of SpicOR29 to limonene enantiomers

To understand the molecular basis of the difference in sensitivity to limonene enantiomers observed for the three OR29 orthologs, we first mapped the amino acid substitutions that distinguish SpicOR29 from its orthologs, SlituOR29 and SlitOR29. We identified eight substitutions, all located in transmembrane helices: TM1 (F39L, I47V), TM2 (L72M), TM3 (F131L), TM4 (T209A, A216V), TM5 (S289C), and TM7b (S379P) (Fig. 4a). Fpocket cavity detection (Le Guilloux et al. 2009), coupled with structural alignment of AlphaFold2 models to published ligand-bound cryo-EM structures, revealed that the I47V substitution is the only one that resides among the 19 residues predicted to constitute the orthosteric binding region (Fig. 4a, b).

**Figure 4:**
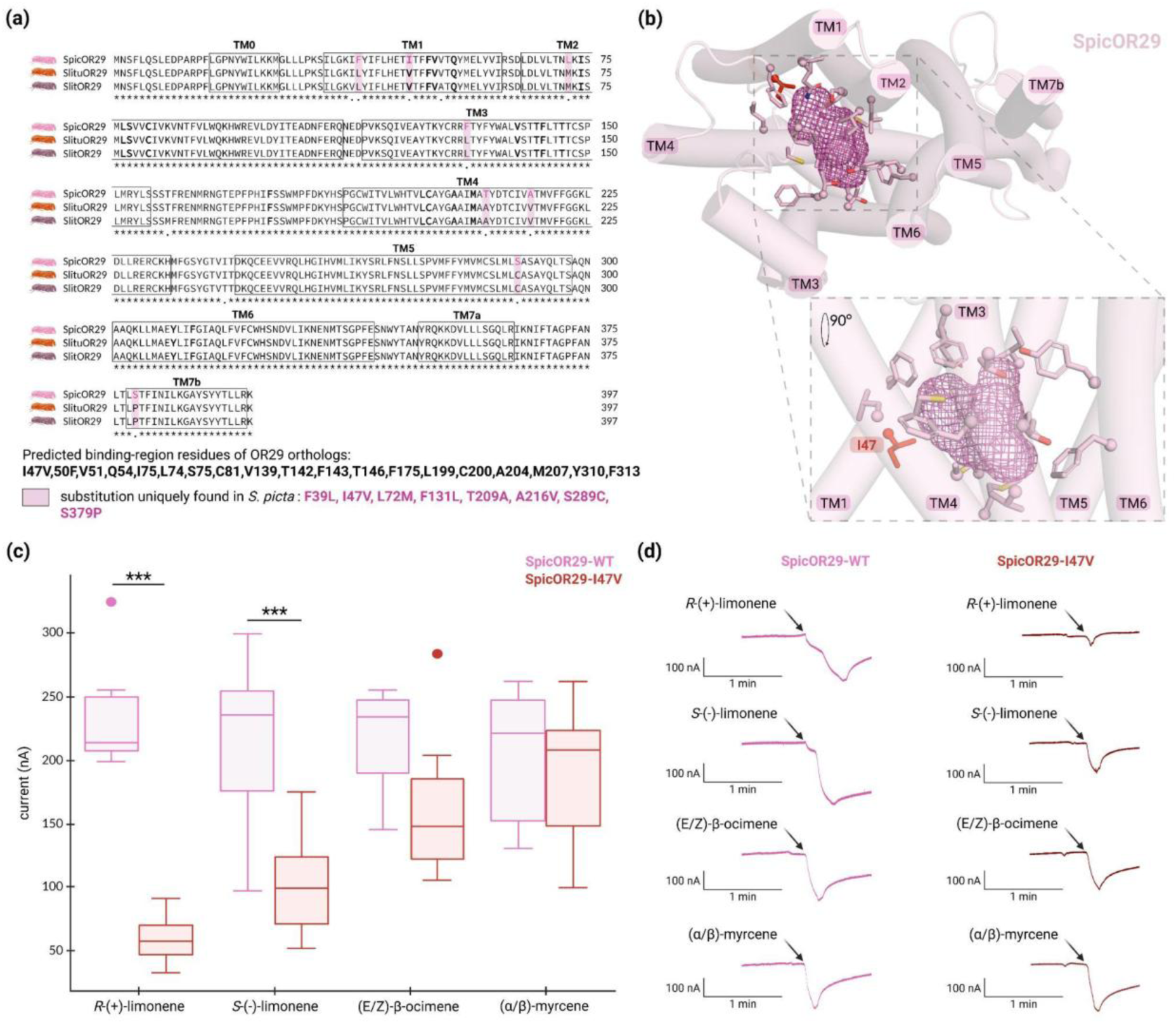
Directed mutagenesis revealed that a single residue in SpicOR29 is crucial for increased sensitivity to limonene enantiomers compared to its orthologs. (**a**) Sequence alignment of OR29 orthologs from *S. picta* and the two polyphagous sister species *S. litura* and *S. littoralis*. The positions of the transmembrane domains (TMs) are indicated by black boxes. Residues predicted to form the binding region are shown in bold. Substitutions unique to SpicOR29 are highlighted in pink. (**b**) Top view of the SpicOR29 AlphaFold2 model, with an inset showing a zoom of the predicted binding region. The receptor cavity is represented as a purple mesh. Side chains of residues predicted to contribute to the binding region are shown as sticks, and the residue mutated in d) is highlighted in red. This figure was generated using the molecular visualization software PyMol (DeLano 2002). (**c**) Box plots showing responses of *Xenopus* oocytes co-expressing SlitOrco with the wild-type SpicOR29 (SpicOR29-WT; light pink) or the single-point mutant (SpicOR29-I47V; red) (n = 8–11) to a 0.1 µM dilution of compounds. The whiskers are extended to 1.5 times the interquartile range from the quartiles.

We thus generated a single-point mutant of SpicOR29, SpicOR29-I47V, in which isoleucine at position 47 was replaced with the valine present in both SlitOR29 and SlituOR29, and compared its function to wild-type SpicOR29 (SpicOR29-WT). This functional study was performed using two-electrode voltage-clamp recordings in *Xenopus laevis* oocytes, rather than single-sensillum recordings from OSNs of transgenic flies, as previously used to functionally characterize Or29 orthologs. This approach was adopted given the time required to generate and establish the corresponding transgenic fly line. Both constructs were tested with a panel of four compounds, including the two limonene enantiomers and two positive controls that elicited comparable sensitivities across all three OR29 orthologs in single-sensillum recordings, namely (α/β)-myrcene and (E/Z)-β-ocimene (Fig. 3a. Consistent with observations in *Drosophila* ab3A OSNs, SpicOR29-WT in the *Xenopus* oocyte system was activated by *R*-(+)-limonene, *S*-(−)-limonene, (α/β)-myrcene, and (*E*/*Z*)-β-ocimene (Fig. 3a,Fig. 4c and d). At the used concentration of 0.1 µM, SpicOR29-I47V exhibited a significantly reduced response to both *S*-(−)-limonene and *R*-(+)-limonene compared with SpicOR29-WT, while its responses to (α/β)-myrcene and (*E*/*Z*)-β-ocimene remained comparable (Fig. 4c and d). These results showed that the single-mutation at position 47 was sufficient to decrease the response of SpicOR29 to limonene enantiomers.

Outliers are indicated by dots. Asterisks indicate statistically significant differences between responses of oocytes expressing different ORs to the same compound (linear model: response ∼ gene × compound, followed by multiple-comparison post hoc tests with a Bonferroni correction; ***, padj < 0.001). (**d**) Representative traces obtained by recording *Xenopus* oocytes expressing SpicOR29-WT (light pink) or the SpicOR29-I47V (red) stimulated with increasing doses of *R*-(+)-limonene. The onset of stimulation is indicated by a black arrow. Black scale bars indicate time and voltage.

## Discussion

The lily moth *S. picta* has evolved a narrow host plant range restricted to certain Amaryllidaceae, whereas two sister species, *S. littoralis* and *S. litura*, are highly polyphagous. In this work, we focused on ORs to investigate if chemosensation may diverge between these three species, as a possible driver of *S. picta* specialization. While major OR expansion or contraction events often reflect species adaptation to ecological niches (Robertson et al. 2019), we did not evidence any such event in the *S. picta* OR repertoire that we manually annotated in the available genome (Wang et al. 2024a). The *S. picta* OR repertoire was comparable in size and composition to those of the two sister species, *S. littoralis* and *S. litura*. We identified only four putative OR pseudogenes in the *S. picta* genome (SpicOR30, 38, 72, and 76), suggesting minimal gene loss despite host specialization. Although pseudogenization does not always result in loss of function in insect chemoreceptors (Prieto-Godino et al. 2016), none of these ORs were detected in the *S. picta* adult antennal transcriptome assembled in this study, suggesting they are unlikely to play an ecological role at this life stage. Limited variation in OR repertoire size despite contrasting host ranges has also been reported across diverse taxa. In *Bombus* bees and *Drosophila* flies, host specialization was associated with relatively small reductions in OR repertoire size (8% and 11%, respectively), whereas in *Papilio* butterflies, specialization was accompanied by a 6% increase (Yin et al. 2022; Singh et al. 2025; McBride 2007). These observations suggest that host specialization does not necessarily drive extensive turnover in OR gene number, especially when speciation is recent. To explain the limited variation in the *S. picta* OR repertoire, we propose two hypotheses that may coexist: (1) *S. picta* specialization may be evolutionarily recent, leaving insufficient time for major gene loss or gain, or (2) strong selection has retained OR diversity, potentially due to ecological pressures. While only a subset of ORs may be sufficient to identify volatiles produced by Amaryllidaceae inflorescences, the remaining ORs could serve to discriminate against non-host plants or mediate recognition of other organisms, such as conspecifics at different life stages, or predators.

Adaptation of the OR repertoire in insect peripheral olfaction involves not only changes in gene number, such as contractions or expansions, but also variation in expression and/or functional divergence of receptors. In moths, a single or a few amino acid changes can alter OR tuning, as demonstrated for pheromone receptors in *Ostrinia* and *Spodoptera* (Leary et al. 2012; Li et al. 2023). To investigate potential substitutions affecting receptor function, we developed a structure-based approach that predicts functional divergence among orthologs. Using large-scale virtual screening, we generated “interaction fingerprints” for each OR, mapping each receptor’s predicted ligand-binding spectrum. In total, we screened 120,591 molecules against 213 ORs from *S. picta*, *S. litura*, and *S. littoralis*. Unsupervised clustering of the profiles revealed that, of the 69 ORs with identified 1:1:1 orthologs across *S. picta*, *S. litura*, and *S. littoralis*, 24 exhibited ligand-binding profiles that occupied distinct regions of the chemical space depending on the ortholog, despite high sequence similarity. By integrating this functional divergence distance with OR expression levels, we identified OR29 as a candidate of interest. This receptor exhibited a markedly divergent predicted ligand-binding profile in *S. picta* compared to its orthologs in *S. litura* and *S. littoralis*, and showed high expression in *S. picta* adult antennae. With 98% and 97% amino acid identity to SlituOR29 and SlitOR29, respectively, SpicOR29 would likely be overlooked by classical sequence-based approaches for detecting functional divergence among orthologs. While our original approach has inherent limitations, including uncertainties in model accuracy, conformational flexibility, and docking precision, as revealed by some predicted binders being not experimentally active and vice versa (Fig. 3a), it offers a rapid and broadly applicable method for identifying putative functionally divergent orthologs across entire OR repertoires.

Functional characterization of OR29 orthologs revealed that SpicOR29, SlituOR29 and SlitOR29 were broadly tuned receptors responsive to common floral volatiles. Consistent with predictions from our structure-based approach, functional divergence among the orthologs was evident in both specificity and sensitivity. The most striking difference between SpicOR29 and its orthologs in *S. litura* and *S. littoralis* was its markedly higher sensitivity to the limonene enantiomers. To gain further molecular insight into the mechanisms behind this functional divergence, we compared the protein sequences of the three OR29 orthologs, mapped the divergent amino acids onto the SpicOR29 3D model, and conducted directed mutagenesis. Only one of the eight amino acid substitutions (I47) between SpicOR29 and its orthologs in *S. litura* and *S. littoralis* was located within the predicted binding region in TM1 and appeared as the most promising residue for directed mutagenesis. Notably, analogous positions in TM1 have been identified as critical determinants of odorant recognition in other *Spodoptera* ORs. For instance, recognition of nonanal by OR47 orthologs in *S. frugiperda* and *S. litura* requires two conserved TM1 residues, H57 and E61 (Wang et al. 2023a). We substituted I47 in SpicOR29 with the corresponding valine found in SlituOR29 and SlitOR29 (I47V) and found that this single change was sufficient to significantly alter the receptor’s recognition of both R-(+)-limonene and S-(−)-limonene, without affecting the recognition of other SpicOR29 ligands, such as (α/β)-myrcene and (E/Z)-β-ocimene. Thus, SpicOR29 single mutant successfully recapitulated the functional properties of OR29 orthologs in *S. litura* and *S. littoralis*, confirming the central role of the binding region in odorant recognition, as reported in previous studies (Leary et al. 2012; Yuvaraj et al. 2021; Li et al. 2023; Wang et al. 2023a; Xu et al. 2023; Ma et al. 2024; Wang et al. 2024b; Zhao et al. 2024).

Limonene enantiomers have been detected in the headspace of inflorescences of certain *S. picta* host plants, such as *C. miniata* (Kiepiel and Johnson 2014), but as not yet been reported in *C. asiaticum* (Miyake et al. 1998). However, these compounds have been identified in other *Crinum* species (Manning and Snijman 2002; Báez et al. 2011; Supplementary Fig. S1a), raising the possibility that their apparent absence from *C. asiaticum* reflects methodological limitations rather than a true absence. Indeed, the failure to detect limonene enantiomers in the study by Miyake *et al*. may be attributable, at least in part, to their dynamic headspace protocol (drawing an excessive 288 L of air through just 50 mg of Tenax TA with solvent elution), which causes breakthrough and evaporation losses of highly volatile compounds such as limonene. In addition, the analytical sensitivity of the GC–MS instrumentation available at the time may have further limited their detection. We therefore re-examined the floral headspace of three *C. asiaticum* inflorescences using 45-min static solid-phase microextraction (SPME) protocol with direct thermal desorption and high-sensitivity GC–MS analysis. We confirmed the presence of limonene enantiomers, demonstrating that these compounds are indeed emitted by *C. asiaticum* inflorescences. Together with the linalool enantiomers, limonene enantiomers are thus the only odorants consistently detected across all examined *S. picta* host inflorescences (Supplementary Fig. S1).

(R/S)-limonene, like other host plant volatiles detected by SpicOR29, does not uniquely characterize the Amaryllidaceae but is instead widespread across angiosperm floral emissions. However, such chemical ubiquity does not preclude ecological relevance, as herbivorous insects often discriminate host plants based on specific combinations and relative ratios of common volatiles (Bruce and Pickett 2011). Interestingly, the presence or absence of limonene enantiomers in synthetic blends mimicking the headspace of cotton, a host plant of *S. littoralis*, influenced the oriented flight of females towards the odor source (Borrero-Echeverry et al. 2015). We propose that *S. picta* may exploit a blend of floral odorants in specific ratios as a family-level olfactory signature for host identification, with limonene enantiomers potentially serving as key components, as reported for its sister species *S. littoralis*. The high expression and enhanced sensitivity of SpicOR29 may enable *S. picta* moths to detect these compounds with exceptional precision.

The expression pattern of SpicOR29 suggests that this receptor may be involved in multiple behavioral processes related to host selection. It is among the most highly expressed olfactory receptors in female antennae, which could suggest a potential role in oviposition site selection. Its notable expression in male antennae raises the possibility that males might also rely on host plant cues to locate females (Xu and Turlings 2018). In addition, expression in both sexes may reflect a role in locating nectar sources for adult feeding. Further studies could clarify the functional significance of SpicOR29 in feeding, mate localization, and oviposition behavior in *S. picta*. Future studies could also investigate the presence and potential role of SpicOR29 in larvae, as we observed strong larval attraction to inflorescences during field experiments in Australia (supplementary fig. S2).

Although we focused on OR29 in this study, it is unlikely to be the only OR involved in *S. picta* host specialization. In *S. littoralis,* floral compounds such as linalool enantiomers and 3-carene are detected by multiple receptors (de Fouchier et al. 2017), highlighting the combinatorial nature of olfactory coding in host recognition. Our structure-based approach predicted functional differences in 24 additional SpicORs when compared to *S. litura* and *S. littoralis.* Further characterization of these receptors may reveal additional changes in ligand specificity or sensitivity, similar to those observed in OR29 lineage, potentially supporting a role for the studied lily volatiles in the host specialization of S. picta. The expression profile of these new candidates should be investigated not only in the antennae but also in the maxillary palps and proboscis, as ORs expressed in mouthparts have been implicated in host finding and food search in other insects, including *Scaptomyza* and *Drosophila* (Matsunaga et al. 2025; Dweck et al. 2016). Moreover, the pseudogenization of four ORs in *S. picta* may have reduced the breadth of volatile compounds detected by this species, compared to the two sister species with broader host ranges. It is also likely that other classes of chemosensory genes contribute to host plant recognition in *S. picta*. Notably, gustatory receptors (GRs), involved in contact chemodetection, and ionotropic receptors (IRs), involved in both olfaction and taste, have been shown to play key roles in host specialization in insects such as *Bombyx mori* and *Drosophila sechellia* (Prieto-Godino et al. 2017; Zhang et al. 2019). Importantly, our study focused on the peripheral olfactory system, whereas host specialization is known to involve modifications at multiple levels, including both peripheral and central neural circuits. In *D. sechellia*, for instance, host adaptation has been associated with rewiring of central olfactory connections (Dürr et al. 2025).

## Conclusion

By integrating genomic and transcriptomic data with an innovative structure-based approach, we identified SpicOR29, a broadly tuned receptor highly expressed in both male and female *S. picta* adult antennae as a potential driver of *S. picta* host plant specialization. We revealed that this receptor has functionally diverged from its orthologs in *S. litura* and *S. littoralis*: during the host specialization of *S. picta*, it has acquired heightened sensitivity to limonene enantiomers, key volatiles of its Amaryllidaceae host plants. Mutagenesis experiments pinpointed one critical amino acid within the predicted binding region that underlies this functional divergence. This case exemplifies how subtle molecular changes, such as a single amino acid substitution, can reshape receptor sensitivity and possibly behavioral outcomes, without requiring large-scale changes in gene number. Future investigations in other narrowly specialized species, such as *S. pectinicornis*, will be essential to determine whether similar mechanisms have been repeatedly selected during evolution.

## Materials and Methods

### OR gene family annotation in *S. picta* and orthology analysis with sister species

A dataset of manually annotated OR amino acid sequences from the reference genome assemblies available at NCBI for *S. exigua* (GenBank assembly GCA_902829305.4), *S. frugiperda* (GCA_023101765.3), *S. litura* (GCA_002706865.3) and *S. littoralis* (GCA_902850265.1) was used as a query to identify ORs in the *S. picta* published genome (GCA_038387885.1; Wang et al. 2024a). The search was performed using TBLASTN 2.14.1 with an e-value cutoff of 1e-04 (Cock et al. 2015).

Changes in the OR repertoire size (expansion or contraction) between the three sister species *S. littoralis*, *S. litura* and *S. picta* were analyzed through phylogenetic reconstruction. OR amino acid sequences were aligned using the MAFFT 7 web server with default settings (Katoh et al. 2019). A phylogenetic tree was generated using the PhyML 3.0 (Guindon et al. 2005) webserver and visualized with FigTree 1.4.4 (https://github.com/rambaut/figtree/releases/tag/v1.4.4). Branch support was determined by performing 100 bootstrap iterations.

### Repertoire-wide virtual screening of odorant receptors in *S. picta*, *S. litura*, and *S. littoralis*

#### Chemical library and receptor preparation

The protocols used to prepare both the chemical library and the receptors followed previously established procedures (Comte et al., 2025). A total of 120,591 molecules, predicted to be volatile, were selected from the COCONUT online database (latest updates: January 2022; Sorokina et al. 2021) for virtual screening on ORs. Molecules were screened against relaxed OR models generated with AlphaFold2 (Jumper et al. 2021) for 213 of the 221 ORs from *S. picta*, *S. litura* and *S. littoralis*. The sequences used to model the 3D structures of all the receptors were obtained from the reference genome assemblies available at NCBI (GenBank accessions GCA_002706865.3, GCA_902850265.1 and GCA_038387885.1). Incomplete sequences (SpicOR22, SlituOR44, SlituOR51, SlituOR64) and predicted pseudogenes (SpicOR30, SpicOR38, SpicOR72, SpicOR76) were excluded from the analysis. The pLDDT score was analyzed across the entire sequence and the binding region of each receptor to validate the accuracy of the selected models (supplementary table S2). The OR 3D models were obtained prior to the release of the latest AlphaFold version (Abramson et al. 2024). However, no significant difference was observed in the modeling of SpicORs, SlituORs and SlitORs between the two versions of AlphaFold (supplementary table S2).

#### Docking simulations

The binding areas of SlitOR25 and SlitOR31 were previously defined (Comte et al. 2025) using available cryo-EM structures of insect ORs (del Mármol et al. 2021; Wang et al. 2024b; Zhao et al. 2024) and Fpocket 4.0 (Le Guilloux et al. 2009). Models of all SpicORs, SlituORs and SlitORs were structurally aligned with the models of SlitOR25 and SlitOR31 using PyMOL 2.5.4 (DeLano 2002). The grid box used for docking simulations was the same as that defined for the screening of SlitOR25 and SlitOR31 (center_x = 10, center_y = -1.67, center_z = -12, size_x = 13, size_y = 13.9, size_z = 15) (Comte et al. 2025). The chemical library was docked using the Vinardo scoring function (Quiroga and Villarreal 2016), implemented in the smina software (Oct 15, 2019, based on AutoDock Vina 1.1.2), weighted by the number of heavy atoms (Comte et al. 2025).

#### Analysis of the predicted chemical space detected by SpicORs, SlituORs and SlitORs

A vector of 120,591 dimensions, containing the re-evaluated Vinardo docking scores for all docked molecules, was generated for each SpicOR, SlituOR and SlitOR. Principal Component Analysis (PCA) was applied to reduce the 120,591-dimensional vectors to 113 dimensions while retaining 99% of the variance. Data clustering was then performed using HDBSCAN 0.8.38 (Campello et al. 2013), with a minimum cluster size of 2, Euclidean distance and default parameters. For illustrative purposes, UMAP 0.5.4 (McInnes et al. 2018) was applied to project the 120,591-dimensional vectors of each SpicOR, SlituOR and SlitOR into two dimensions using *n_neighbors* = 10, *min_dist* = 0.0, and default settings. HDBSCAN clustering (Campello et al. 2013) was then performed on the reduced representation to visually highlight distinct groups based on the predicted chemical space.

We ranked 120,591 molecules screened against SpicOR29, SlituOR29, and SlitOR39 by their weighted docking scores to predict the chemical space recognized by OR29 orthologs. The lowest 6% of scoring compounds were identified as potential binders, in accordance with the methodology described by Comte et al. (2025).

### RNA-seq analyses

#### Collection and rearing of *Spodoptera picta*

Collection sites around Sydney, Australia, were identified through user posts on the online social network iNaturalist (Matheson CA 2014). The presence of *S. picta* and its primary host plant in the region, *C. pedunculatum* (Ang et al. 2010), was checked between 2012 and 2022 (supplementary figure S2). Larvae at various developmental stages were collected on *C. pedunculatum*, at coordinates 34°19’35.8”S, 150°55’16.9”E (5 Hamilton Rd, Thirroul, NSW 2515, Australia). The larvae were reared in plastic containers at 25°C and supplied with fresh *C. pedunculatum* tissues, primarily inflorescences, every two days until they reached the imago stage.

#### Sample preparation, RNA extraction and sequencing

Antennae from virgin adult males (n=9) and virgin adult females (n=23) were collected two to three days post-emergence, pooled into three biological replicates per sex and preserved in RNA*later*™ (Thermo Fisher Scientific, Waltham, MA, USA) for shipment to France (Export permit from the Australian Government PWS2022-AU-002455). Upon arrival in France, samples were transferred to TRIzol™ Reagent (Thermo Fisher Scientific). Tissues were homogenized using a Polytron™ Pro 200 (Pro Scientific Inc., Oxford, CT, USA), followed by phenol-chloroform RNA extraction. RNA quality (260/280 nm ratio) and quantity were assessed using a NanoDrop™ ND-2000 spectrophotometer (Thermo Fisher Scientific). The six RNA samples were shipped on dry ice to Novogene Company Limited (Cambridge, UK) for directional library construction and sequencing. Sequencing was performed on an Illumina NovaSeq 6000 platform with a paired-end read length of 150 bp, generating 18 Gb of clean data per library.

#### Transcriptome assembly and differential expression analysis

The Galaxy server hosted by the BioInformatics Platform for Agro-ecosystem Arthropods (Rennes, France) was used for data processing and analysis. Raw read quality was assessed using FastQC 0.74 (Andrews 2010), and low-quality reads were trimmed with Trimmomatic 0.39 (Bolger et al. 2014) using the following parameters: sliding window = 4, average quality ≥ 30, minimum length = 20 bases, and headcrop = 10 bases. De novo assembly was performed with Trinity 2.8.5 using default parameters (Grabherr et al. 2011), and transcriptome completeness was evaluated using BUSCO 5.5.0 (Simão et al. 2015). Coding sequences were extracted from the reference transcriptome using TransDecoder v5.5.0 (Haas et al. 2013), setting a minimum protein length of 50 amino acids. Redundant sequences were subsequently clustered with CD-HIT 4.8.1 (Fu et al. 2012), using a similarity threshold of 0.95 and a word size of 10.

The newly annotated SpicORs sequences, derived from the genome published by Wang et al., 2024a (GenBank accession GCA_038387885.1), were used as queries to identify corresponding transcripts in the *S. picta* transcriptome assembly. This was done using TBLASTN 2.14.1 (Cock et al. 2015), with an E-value cutoff of 1e−04. Transcript counts and TPM values for *S. picta* samples were obtained by mapping clean reads to the OR transcript sequences using Kallisto quant 0.48.0 (Bray et al. 2016). Differential expression of transcripts between sexes was calculated using DESeq2 2.11.40.8 (Love et al. 2014). A gene was considered differentially expressed if the adjusted p-value (Benjamini-Hochberg correction) was below 0.1.

### Functional study of SpicOR29, SlituOR29 and SlitOR29

#### Chemical panel selection

The compound panel used in the electrophysiological assays comprised 9 of the 12 molecules reported as constituents of the floral headspace of C. asiaticum and C. miniata (Miyake et al. 1998; Kiepiel and Johnson 2014; see Supplementary Fig. S1a), selected on the basis of our SBVS analysis, which predicted binding to at least one OR29 ortholog. The panel was complemented with previously characterized SlitOR29 ligands that were also predicted to bind SpicOR29 or SlituOR29 (de Fouchier et al. 2017). Both enantiomers of limonene were tested. Compounds were obtained from AMBINTER (Orléans, France). Depending on their solubility, compounds were either directly mixed with paraffin oil or introduced into a DMSO stock solution before dilution in paraffin oil. We made sure that the final concentration of DMSO did not exceed 5%.

#### Heterologous expression in *Drosophila melanogaster* ab3A olfactory neurons

The open reading frames of SpicOR29, SlituOR29 and SlitOR29 were synthesized in vitro by Synbio Technologies (Monmouth Junction, NJ, USA) and subcloned into the pUAST.attB vector. Sequences were codon-optimized for expression in *Drosophila melanogaster*. The pUAST.attB-SpicOR29, pUAST.attB-SlituOR29 and pUAST.attB-SlitOR29 plasmids were injected into *D. melanogaster* embryos (genotype y1 M{vas-int.Dm}ZH-2A w*; M{3xP3-RFP.attP}ZH-86Fb) by BestGene Inc. (Chino Hills, CA, USA), utilizing the φC31 integrase system. This facilitated the insertion of the constructs into the genomic locus 86Fb on the third chromosome. UAS-SpicOR29, UAS-SlituOR29 and UAS-SlitOR29 lines were then crossed with Df(2L)Or22ab, TI{GAL4}Or22ab stock flies to generate homozygous flies expressing the OR of interest in the *D. melanogaster* ab3 sensilla, replacing OR22a and OR22b. Transgenic flies were reared on a standard cornmeal-yeast-agar medium and maintained in a Sanyo incubator (MIR-553) at 25°C under a 12-hour light-dark photoperiod.

#### Single-sensillum recordings

Transgenic flies were transferred to 29°C 24 hours prior to single-sensillum recordings to optimize GAL4 activity while minimizing any impact on line viability (Duffy 2002). Male and female flies, aged 5 days, were randomly selected from the *Drosophila* population. Single-sensillum recordings were conducted following established protocols (Comte et al. 2025).

The odorant stimuli for the screening experiment were prepared by loading 10 µl of each odorant solution (10 µg/µl) onto a 1 cm² filter paper and placing it into a Pasteur pipette. Dose-response analyses were performed using doses ranging from 0.001 µg to 100 µg in the stimulus cartridges. Neuronal activity was recorded and analyzed using pCLAMP 10. Net responses of ab3A neurons were determined by subtracting the spontaneous firing rate from the firing rate during the odorant stimulation, as described previously (de Fouchier et al. 2017). Three diagnostic stimuli were used to distinguish ab3 sensilla from other basiconic sensilla: 10 µg of ethyl 3-hydroxybutyrate, 10 µg of ethyl acetate and CO_2_ from human expiration. These stimuli activate ab3B OSNs, ab2A OSNs and ab1C OSNs, respectively (Münch and Galizia 2016). The absence of the endogenous receptor OR22a in the ab3A neuron was confirmed by delivering 0.1 µg of ethyl hexanoate, a potent ligand for DmelOR22a (Münch and Galizia 2016).

Eight biological replicates were performed for each compound and dose tested. Odorants were considered as active if they elicited a neuronal response at 100 µg that was significantly different from that induced by the solvent control during the screening experiment (Friedman test, followed by Dunn’s test with Benjamini & Yekutieli correction, p < 0.05). For dose–response experiments, data were fitted with sigmoidal curves using the curve_fit function from the SciPy 1.16.0 library, and EC₅₀ values were estimated from the fitted curves. Statistical analyses were performed using R 4.3.2 and Python 3.11.5.

#### Site-Directed Mutagenesis of SpicOR29

The functional characterization of both the mutant and wild-type receptors was conducted using the two-electrode voltage clamp system as previously described (Cao et al. 2022). The open reading frames of the ORs were synthesized by Synbio Technologies and subcloned into the pCS2+ vector. The resulting plasmids were linearized with NotI and used as templates for *in vitro* transcription of capped cRNAs with SP6 RNA polymerase, using the HiScribe SP6 RNA Synthesis Kit (New England Biolabs, Ipswich, MA, USA) according to the manufacturer’s instructions. A mixture of 27.6 ng of SlitOrco cRNA and 27.6 ng of SpicOR29 cRNA (wild-type or site-directed mutant) was microinjected into stage V-VII *Xenopus* oocytes using a Nanoject III microinjector (Drummond Scientific Company, Broomall, PA, USA). After 3-4 days of incubation at 18℃, the response of individual oocytes was recorded using a two-electrode voltage clamp set-up (TURBO-TEC-03X, npi electronic GmbH, Tamm, Germany) at a holding potential of -80 mV. Data acquisition and analysis were performed with a Digidata 1550 and pCLAMP 10.6 software (Axon Instruments Inc., Union City, CA, USA). Each oocyte expressing either the mutant or wild-type OR was tested with a single compound. Seven biological replicates were recorded for each compound tested on the wild-type OR, and eleven replicates for the mutant OR. Statistical analyses were performed using R 4.3.2. A linear model was fitted to the data, and multiple comparisons were adjusted using the Bonferroni correction with the emmeans 1.11.2-8 package.

Stock solutions of the test compounds were prepared in DMSO at 1 mol/L and subsequently diluted with 1 × Ringer’s buffer (96 mM NaCl, 2 mM KCl, 5 mM MgCl_2_, 0.8 mM CaCl_2_, and 5 mM HEPES, pH 7.6) to a final concentration of 10^-4^ M for the scanning experiments. 1 × Ringer’s buffer containing 0.1% DMSO served as a negative control.

#### Headspace sampling and GC–MS analysis of *Crinum asiaticum* inflorescences

Limonene enantiomers have not been previously reported in the inflorescence headspace of *C. asiaticum* (Miyake et al. 1998) but have been identified in two other *Crinum* species (Manning and Snijman 2002; Báez et al. 2011; see supplementary figure S1a). To check the presence of limonene enantiomers in *C. asiaticum*, static sampling was performed using solid-phase microextraction (SPME) on three inflorescences of *C. asiaticum* in a greenhouse at Park Phoenix (Nice, France). The inflorescences of *C. asiaticum* were isolated inside a one-meter Nalophan bag, which was sealed with clamps. A hole was made in the bag with a needle to insert a 50/30 µm DVB/CAR/PDMS stableflex fiber for SPME. The fiber was exposed inside the bag for 45 minutes. Headspace extraction analyses were performed using an Agilent 8890 gas chromatograph (GC) coupled with an Agilent 5977B single quadrupole Mass Spectrometry Detector. The capillary column used was an HP-5 MS column (30 m length × 0.25 mm internal diameter × 0.25 μm film thickness; Agilent Technologies). Helium was employed as the carrier gas at a constant flow rate of 1.0 mL/min. The oven temperature program began at 50°C, with a heating rate of 8°C/min until reaching 300°C. Ionization was carried out in electron impact (EI) mode at 70 eV, with full scan mode used over a range of 30 to 400 m/z. The source temperature was set to 230°C, and the quadrupole temperature was maintained at 150°C. GC/MS data were analyzed using Agilent MassHunter software, and compound identification was achieved by comparing the spectra with the NIST 2020 library and matching the retention index with commercially available analytical standards (Sigma-Aldrich). Pure standards of limonene enantiomers were run under identical conditions to confirm compound identities (supplementary figure S1b).

## Acknowledgements

The authors are grateful to the Université Côte d’Azur’s Center for High-Performance Computing (OPAL infrastructure) for providing resources and support. We thank the staff of the Parc Phoenix, Nice, France, for allowing us to perform the floral headspace collection of *Crinum asiaticum* in the Parc.

## Data availability

RNAseq raw data sequences from male and female *S. picta* antennae have been deposited at the National Center for Biotechnology Information - Sequence Read Archive database under accession numbers (pending).

## Contributions

EJJ and SF designed research; AC, ML, SZ, AT, RM and JG performed research; AC analyzed data; AC, ML, EJJ and SF wrote the paper. All authors discussed the results, reviewed and approved the manuscript.

## References

Abramson J, Adler J, Dunger J, Evans R, Green T, Pritzel A, Olaf Ronneberger O, Willmore L, Ballard AJ, Bambrick J et al. 2024. Accurate structure prediction of biomolecular interactions with AlphaFold 3. Nature 630(8016):493–500. 10.1038/s41586-024-07487-w.

Álvarez-Ocaña R, Shahandeh MP, Ray V, Auer TO, Gompel N, Benton R. 2023. Odor-regulated oviposition behavior in an ecological specialist. Nature Communications 14(1):3041. 10.1038/s41467-023-38722-z.

Andersson MN, Löfstedt C, Newcomb RD. 2015. Insect olfaction and the evolution of receptor tuning. Frontiers in ecology and evolution 3:53. 10.3389/fevo.2015.00053.

Andersson MN, Keeling CI, Mitchell RF. 2019. Genomic content of chemosensory genes correlates with host range in wood-boring beetles (*Dendroctonus ponderosae*, *Agrilus planipennis*, and *Anoplophora glabripennis*). BMC genomics 20(1):690. 10.1186/s12864-019-6054-x.

Andrews S. 2010. FastQC: a quality control tool for high throughput sequence data. http://www.bioinformatics.babraham.ac.uk/projects/fastqc.

Ang WF, Teo S, Lok AFS, Suen SM, Ng BY. 2010. Late instar caterpillar and metamorphosis of *Spodoptera picta* Guérin-Méneville (Lepidoptera: Noctuidae: Noctuinae) with notes on its cannibalistic behaviour. Nat. Singapore 3:239–244.

Auer TO, Khallaf MA, Silbering AF, Zappia G, Ellis K, Álvarez-Ocaña R, Arguello JR, Hansson BS, Jefferis GSXE, Caron SJC et al. 2020. Olfactory receptor and circuit evolution promote host specialization. Nature 579(7799):402–408. 10.1038/s41586-020-2073-7.

Báez D, Pino JA, Morales D. 2011. Scent Composition in Some Cuban Flowers: *Clitoria Fairchildiana* RA Howard, *Brunfelsia Nitida* Benth. and *Crinum Oliganthum* Urban. Journal of Essential Oil Bearing Plants 14(4):383–386. 10.1080/0972060X.2011.10643590.

Bauder JAS, Karolyi F. 2019. Superlong proboscises as co-adaptations to flowers. In: Krenn HW, editor. Insect Mouthparts: Form, Function, Development and Performance. Cham: Springer, Charm. p. 479–527. 10.1007/978-3-030-29654-4_15.

Bolger AM, Lohse M, Usadel B. 2014. Trimmomatic: a flexible trimmer for Illumina sequence data. Bioinformatics 30(15):2114–2120. 10.1093/bioinformatics/btu170.

Borrero-Echeverry F, Becher PG, Birgersson G, Bengtsson M, Witzgall P, Saveer AM. 2015. Flight attraction of *Spodoptera littoralis* (Lepidoptera, Noctuidae) to cotton headspace and synthetic volatile blends. Frontiers in Ecology and Evolution 3:56. 10.3389/fevo.2015.00056.

Bray NL, Pimentel H, Melsted P, Pachter L. 2016. Near-optimal probabilistic RNA-seq quantification. Nature biotechnology 34(5):525–527.10.1038/nbt.3519.

Bruce TJ, Pickett JA. 2011. Perception of plant volatile blends by herbivorous insects–finding the right mix. Phytochemistry 72(13):1605–1611. 10.1016/j.phytochem.2011.04.011.

Campello RJ, Moulavi D, Sander J. 2013. Density-based clustering based on hierarchical density estimates. In: Pei J, Tseng VS, Cao L, Motoda H, Xu G, editors. Pacific-Asia conference on knowledge discovery and data mining. Berlin, Heidelberg: Springer Berlin Heidelberg .p. 160–172.

Cao S, Liu Y, Wang G. 2022. Protocol to identify ligands of odorant receptors using two-electrode voltage clamp combined with the *Xenopus* oocytes heterologous expression system. STAR protocols 3(2):101249. 10.1016/j.xpro.2022.101249.

Ceja-Navarro JA, Karaoz U, Bill M, Hao Z, White III RA, Arellano A, Ramanculova L, Filley TR, Berry TD, Conrad ME et al. 2019. Gut anatomical properties and microbial functional assembly promote lignocellulose deconstruction and colony subsistence of a wood-feeding beetle. Nature microbiology 4(5):864–875. 10.1038/s41564-019-0384-y.

Chahda JS, Soni N, Sun JS, Ebrahim SA, Weiss BL, Carlson JR. 2019. The molecular and cellular basis of olfactory response to tsetse fly attractants. PLoS genetics 15(3): e1008005. 10.1371/journal.pgen.1008005.

Clyne PJ, Warr CG, Freeman MR, Lessing D, Kim J, Carlson JR. 1999. A novel family of divergent seven-transmembrane proteins: candidate odorant receptors in *Drosophila*. Neuron 22(2):327–338. 10.1016/S0896-6273(00)81093-4.

Cock PJ, Chilton JM, Grüning B, Johnson JE, Soranzo N. 2015. NCBI BLAST+ integrated into Galaxy. Gigascience 4(1):s13742–015. 10.1186/s13742-015-0080-7.

Comte A, Lalis M, Brajon L, Moracci R, Montagné N, Topin J, Jacquin-Joly E, Fiorucci S. 2025. Accelerating ligand discovery for insect odorant receptors. International Journal of Biological Sciences 21(5):2101. 10.7150/ijbs.105648.

del Mármol J, Yedlin MA, Ruta V. 2021. The structural basis of odorant recognition in insect olfactory receptors. Nature 597(7874):126–131. 10.1038/s41586-021-03794-8.

de Fouchier A, Walker III WB, Montagné N, Steiner C, Binyameen M, Schlyter F, Chertemps T, Maria A, François MC, Monsempes et al. 2017. Functional evolution of Lepidoptera olfactory receptors revealed by deorphanization of a moth repertoire. Nature communications 8(1):15709. 10.1038/ncomms15709.

DeLano WL. 2002. Pymol: An open-source molecular graphics tool. CCP4 Newsl. Protein Crystallogr 40(1):82–92.

Dong Z, Wang Y, Tian Y, Guan Z, Bai M, Zhang B, Jia Z, Wu J, Cao S, Gong Z et al. 2026. Structural basis of sex pheromone detection in aphids. Cell Research 36:582–594. 10.1038/s41422-026-01267-z.

Duffy JB. 2002. GAL4 system in Drosophila: a fly geneticist’s Swiss army knife. Genesis 34: 1–15. 10.1002/gene.10150.

Dürr BR, Bertolini E, Takagi S, Pascual J, Abuin L, Lucarelli G, Benton R, Auer TO. 2025. Olfactory projection neuron rewiring in the brain of an ecological specialist. Cell Reports 44(5):115615. 10.1016/j.celrep.2025.115615.

Dweck HK, Ebrahim SA, Khallaf MA, Koenig C, Farhan A, Stieber R, Weißflog J, Svatoš A, Grosse-Wilde E, Knaden M et al. 2016. Olfactory channels associated with the *Drosophila* maxillary palp mediate short-and long-range attraction. Elife 5:e14925. 10.7554/eLife.14925.

Fu L, Niu B, Zhu Z, Wu S, Li W. 2012. CD-HIT: accelerated for clustering the next-generation sequencing data. Bioinformatics 28(23):3150–3152.10.1093/bioinformatics/bts565.

Gao Q, Chess A. 1999. Identification of candidate *Drosophila* olfactory receptors from genomic DNA sequence. Genomics 60(1):31–39. 10.1006/geno.1999.5894.

Gikonyo MW, Ahn SJ, Biondi M, Fritzlar F, Okamura Y, Vogel H, Köllner TG, Şen İ, Hernández-Teixidor D, Lee CF et al. 2024. A radiation of *Psylliodes* flea beetles on Brassicaceae is associated with the evolution of specific detoxification enzymes. Evolution 78(1):127–145. 10.1093/evolut/qpad197.

Goldman-Huertas B, Mitchell RF, Lapoint RT, Faucher CP, Hildebrand JG, Whiteman NK. 2015. Evolution of herbivory in Drosophilidae linked to loss of behaviors, antennal responses, odorant receptors, and ancestral diet. Proceedings of the National Academy of Sciences 112(10):3026–3031. 10.1073/pnas.1424656112.

Grabherr MG, Haas BJ, Yassour M, Levin JZ, Thompson DA, Amit I, Adiconis X, Fan L, Raychowdhury R, Zeng Q et al. 2011. Full-length transcriptome assembly from RNA-Seq data without a reference genome. Nature Biotechnology 29(7):644–652. 10.1038/nbt.1883.

Guindon S, Lethiec F, Duroux P, Gascuel O. 2005. PHYML Online—a web server for fast maximum likelihood-based phylogenetic inference. Nucleic acids research 33(suppl_2):W557–W559. 10.1093/nar/gki352.

Haas BJ, Papanicolaou A, Yassour M, Grabherr M, Blood PD, Bowden J, Couger MB, Eccles D, Li B, Lieber M et al. 2013. De novo transcript sequence reconstruction from RNA-seq using the Trinity platform for reference generation and analysis. Nature protocols 8(8):1494–1512. 10.1038/nprot.2013.084.

He P, Engsontia P, Chen GL, Yin Q, Wang J, Lu X, Zhang YN, Li ZQ, He M. 2018. Molecular characterization and evolution of a chemosensory receptor gene family in three notorious rice planthoppers, *Nilaparvata lugens*, *Sogatella furcifera* and *Laodelphax striatellus*, based on genome and transcriptome analyses. Pest Management Science 74(9):2156–2167. 10.1002/ps.4912.

Jang SS, Mandala S, Jiang H, Zhang X, Cole PA, del Mármol J, unpublished data, https://www.biorxiv.org/content/10.1101/2025.05.30.657074v1.full, last accessed August 2, 2025.

Jumper J, Evans R, Pritzel A, Green T, Figurnov M, Ronneberger O, Tunyasuvunakool K, Bates R, Žídek A, Potapenko A et al. 2021. Highly accurate protein structure prediction with AlphaFold. Nature 596(7873):583–589. 10.1038/s41586-021-03819-2.

Katoh K, Rozewicki J, Yamada KD. 2019. MAFFT online service: multiple sequence alignment, interactive sequence choice and visualization. Briefings in bioinformatics 20(4):1160–1166. 10.1093/bib/bbx108.

Kergoat GJ, Goldstein PZ, Le Ru B, Meagher Jr RL, Zilli A, Mitchell A, Clamens AL, Gimenez S, Barbut J, Nègre N et al. (2021). A novel reference dated phylogeny for the genus *Spodoptera* Guenée (Lepidoptera: Noctuidae: Noctuinae): new insights into the evolution of a pest-rich genus. Molecular phylogenetics and evolution 161:107161. 10.1016/j.ympev.2021.107161.

Kiepiel I, Johnson SD. 2014. Shift from bird to butterfly pollination in *Clivia* (Amaryllidaceae). American Journal of Botany 101(1):190–200. 10.3732/ajb.1300363.

Larsson MC, Domingos AI, Jones WD, Chiappe ME, Amrein H, Vosshall LB. 2004. Or83b encodes a broadly expressed odorant receptor essential for *Drosophila* olfaction. Neuron 43(5):703–714. 10.1016/j.neuron.2004.08.019.

Leary GP, Allen JE, Bunger PL, Luginbill, J,B., Linn Jr CE, Macallister IE, Kavanaugh MP, Wanner, K. W. 2012. Single mutation to a sex pheromone receptor provides adaptive specificity between closely related moth species. Proceedings of the National Academy of Sciences 109(35):14081–14086. 10.1073/pnas.1204661109.

Le Guilloux V, Schmidtke P, Tuffery P. 2009. Fpocket: an open source platform for ligand pocket detection. BMC bioinformatics 10(1):168. 10.1186/1471-2105-10-168.

Li Z, Capoduro R, Bastin–Héline L, Zhang S, Sun D, Lucas P, Dabir-Moghaddam D, François MC, Liu Y, Wang G et al. 2023. A tale of two copies: Evolutionary trajectories of moth pheromone receptors. Proceedings of the National Academy of Sciences 120(20):e2221166120. 10.1073/pnas.2221166120.

Liu XL, Zhang J, Yan Q, Miao CL, Han WK, Hou W, Yang K, Hansson BS, Peng YC, Guo JM et al. 2020. The molecular basis of host selection in a crucifer-specialized moth. Current Biology 30(22): 4476–4482. 10.1016/j.cub.2020.08.047.

Love MI, Huber W, Anders S. 2014. Moderated estimation of fold change and dispersion for RNA-seq data with DESeq2. Genome biology 15(12):550. 10.1186/s13059-014-0550-8.

Ma Y, Yang TT, Ni S, Wang JX, He Y, Si YX, Zhang J, Dong SL, Yan Q. 2024. The odorant receptor recognizing camphor in a camphor tree specialist *Orthaga achatina* (Lepidoptera: Pyralidae). Journal of Agricultural and Food Chemistry 72(5):2689–2696. 10.1021/acs.jafc.3c08877.

Manning JC, Snijman D. 2002. Hawkmoth-pollination in *Crinum variabile* (Amaryllidaceae) and the biogeography of sphingophily in southern African Amaryllidaceae. South African Journal of Botany 68(2):212–216. 10.1016/S0254-6299(15)30422-1.

Matheson CA. 2014. inaturalist. Reference Reviews 28(8):36–38. 10.1108/RR-07-2014-0203.

Matsunaga T, Reisenman CE, Goldman-Huertas B, Rajshekar S, Suzuki HC, Tadres D, Wong J, Louis M, Ramírez SR, Whiteman NK. 2025. Odorant receptors mediating avoidance of toxic mustard oils in *Drosophila melanogaster* are expanded in herbivorous relatives. Molecular Biology and Evolution, 42(9):msaf164. 10.1093/molbev/msaf164.

McBride CS, Baier F, Omondi AB, Spitzer SA, Lutomiah J, Sang R, Ignell R, Vosshall LB. 2014. Evolution of mosquito preference for humans linked to an odorant receptor. Nature 515(7526):222–227. 10.1038/nature13964.

McInnes L, Healy J, Melville J. 2018. UMAP: Uniform Manifold Approximation and Projection. Journal of Open Source Software 3(29):861. 10.21105/joss.00861.

Miyake T, Yamaoka R, Yahara T. 1998. Floral scents of hawkmoth-pollinated flowers in Japan. Journal of Plant Research 111(2):199–205. 10.1007/BF02512170.

Montagné N, Gévar J, Lucas P. 2022. Semiochemicals and communication in insects. In: Fauvergue X, Rusch A., Barret M., Bardin M, Jacquin-Joly E, Malausa T, Lannou C, editors. Extended biocontrol. Dordrecht: Springer Netherlands. p. 173–181. 10.1007/978-94-024-2150-7_15.

Münch D, Galizia CG. 2016. DoOR 2.0-comprehensive mapping of *Drosophila melanogaster* odorant responses. Scientific reports 6(1):21841. 10.1038/srep21841.

Prieto-Godino LL, Rytz R, Bargeton B, Abuin L, Arguello JR, Peraro MD, Benton R. 2016. Olfactory receptor pseudo-pseudogenes. Nature 539(7627):93–97. 10.1038/nature19824.

Prieto-Godino LL, Rytz R, Cruchet S, Bargeton B, Abuin L, Silbering AF, Ruta V, Peraro MD, Benton R. 2017. Evolution of acid-sensing olfactory circuits in drosophilids. Neuron 93(3):661–676. 10.1016/j.neuron.2016.12.024.

Quiroga R, Villarreal MA. 2016. Vinardo: A scoring function based on autodock vina improves scoring, docking, and virtual screening. PloS one 11(5):e0155183. 10.1371/journal.pone.0155183.

Robertson HM, Baits RL, Walden KK, Wada-Katsumata A, Schal C. 2018. Enormous expansion of the chemosensory gene repertoire in the omnivorous German cockroach *Blattella germanica*. Journal of Experimental Zoology Part B: Molecular and Developmental Evolution 330(5):265–278. 10.1002/jez.b.22797.

Robertson HM. 2019. Molecular evolution of the major arthropod chemoreceptor gene families. Annual review of entomology 64(1):227–242. 10.1146/annurev-ento-020117-043322.

Simão FA, Waterhouse RM, Ioannidis P, Kriventseva EV, Zdobnov EM. 2015. BUSCO: assessing genome assembly and annotation completeness with single-copy orthologs. Bioinformatics 31(19):3210–3212. 10.1093/bioinformatics/btv351.

Singh A, Pope NS, López-Uribe MM. 2025. Shifts in bee diet breadths are associated with gene gains and losses and positive selection across olfactory receptors. G3: Genes, Genomes, Genetics 15(8): jkaf105. 10.1093/g3journal/jkaf105.

Sorokina M, Merseburger P, Rajan K, Yirik MA, Steinbeck C. 2021. COCONUT online: collection of open natural products database. Journal of Cheminformatics 13(1):2. 10.1186/s13321-020-00478-9.

Vosshall LB, Amrein H, Morozov PS, Rzhetsky A, Axel R. 1999. A spatial map of olfactory receptor expression in the *Drosophila* antenna. Cell 96(5):725–736. 10.1016/S0092-8674(00)80582-6.

Walker III WB, Roy A, Anderson P, Schlyter F, Hansson BS, Larsson MC. 2019. Transcriptome analysis of gene families involved in chemosensory function in *Spodoptera littoralis* (Lepidoptera: Noctuidae). BMC genomics 20(1):428. 10.1186/s12864-019-5815-x.

Wang JX, Wei ZQ, Chen MD, Yan Q, Zhang J, Dong SL. 2023a. Conserved odorant receptors involved in nonanal-induced female attractive behavior in two *Spodoptera* Species. Journal of agricultural and food chemistry 71(37):13795–13804.10.1021/acs.jafc.3c03265.

Wang J, Wei J, Yi T, Li YY, Xu T, Chen L, Xu H. 2023b. A green leaf volatile,(Z)-3-hexenyl-acetate, mediates differential oviposition by *Spodoptera frugiperda* on maize and rice. BMC biology 21(1):140. 10.6084/m9.figshare.23264939.

Wang H, Song J, Hunt BJ, Zuo K, Zhou H, Hayward A, Li B, Xiao Y, Geng X, Bass C et al. 2024a. UDP-glycosyltransferases act as key determinants of host plant range in generalist and specialist *Spodoptera* species. Proceedings of the National Academy of Sciences 121(19), e2402045121. 10.1073/pnas.2402045121.

Wang Y, Qiu L, Wang B, Guan Z, Dong Z, Zhang J, Cao S, Yang L, Wang B, Gong Z et al. 2024b. Structural basis for odorant recognition of the insect odorant receptor OR-Orco heterocomplex. Science 384(6703):1453–1460. 10.1126/science.adn6881.

Wang J, Yang C, Chang S, Jiao D, Lin J, Yang X, Cai W, Ma D, Ding ZJ, Huang J et al. 2026. Cryo-EM structures of *Drosophila* OR67d–Orco complexes reveal insect pheromone sensing mechanism. Cell Research 36:595–610. 10.1038/s41422-026-01264-2.

Xu H, Turlings TC. 2018. Plant volatiles as mate-finding cues for insects. Trends in Plant Science 23(2):100–111. 10.1016/j.tplants.2017.11.004.

Xu L, Jiang HB, Yu JL, Pan D, Tao Y, Lei Q, Chen Y, Liu Z, Wang JJ. 2023. Two odorant receptors regulate 1-octen-3-ol induced oviposition behavior in the oriental fruit fly. Communications Biology 6(1):176. 10.1038/s42003-023-04551-5.

Yang J, Mo BT, Li GC, Huang LQ, Guo H, Wang CZ. 2024. Identification and functional characterization of chemosensory genes in olfactory and taste organs of *Spodoptera litura* (Lepidoptera: Noctuidae). Insect Science 31(6):1721–1742. 10.1111/1744-7917.13350.

Yin N, Xiao H, Yang A, Wu C, Liu N. 2022. Genome-wide analysis of odorant and gustatory receptors in six *Papilio* butterflies (Lepidoptera: Papilionidae). Insects 13(9):779. 10.3390/insects13090779.

Yuvaraj JK, Roberts RE, Sonntag Y, Hou XQ, Grosse-Wilde E, Machara A, Zhang DD, Hansson BS, Johanson U, Löfstedt C et al. 2021. Putative ligand binding sites of two functionally characterized bark beetle odorant receptors. BMC biology 19(1):16. 10.1186/s12915-020-00946-6.

Zdenek CN, Cardoso FC, Robinson SD, Mercedes RS, Raidjõe ER, Hernandez-Vargas MJ, Jin J, Corzo G, Vetter I, King GF et al. 2024. Venom exaptation and adaptation during the trophic switch to blood-feeding by kissing bugs. Iscience 27(9):110723. 10.1016/j.isci.2024.110723.

Zhang ZJ, Zhang SS, Niu BL, Ji DF, Liu XJ, Li MW, Bai H, Palli SR, Wang CZ, Tan AJ. 2019. A determining factor for insect feeding preference in the silkworm, *Bombyx mori*. PLoS Biology 17(2): e3000162. 10.1371/journal.pbio.3000162

Zhao J, Chen AQ, Ryu J, del Mármol J. 2024. Structural basis of odor sensing by insect heteromeric odorant receptors. Science 384(6703):1460–1467. 10.1126/science.adn6384.

