## Supplementary Files for "Single-point mutation alters odorant receptor sensitivity associated with host plant specialization in *Spodoptera* moths"

**Supplementary Table 1. Of the 75 newly annotated genes in the published *S. picta* genome**
**(GenBank accession GCA\_038387885.1; Wang et al. 2024a), 62 were identified in the *S. picta***
**adult antennal transcriptome.** In the table, complete genes are highlighted in green, incomplete genes
in red, and pseudogenes in orange, while SpicORs absent from the transcriptome are indicated in grey.
Sequences used for SpicOR modeling were taken from genome annotations, except for SpicOR22,
which was obtained from the antennal transcriptome (truncated in the genome).
<u>table S1 is provided in the Table\_S1\_xls file</u>

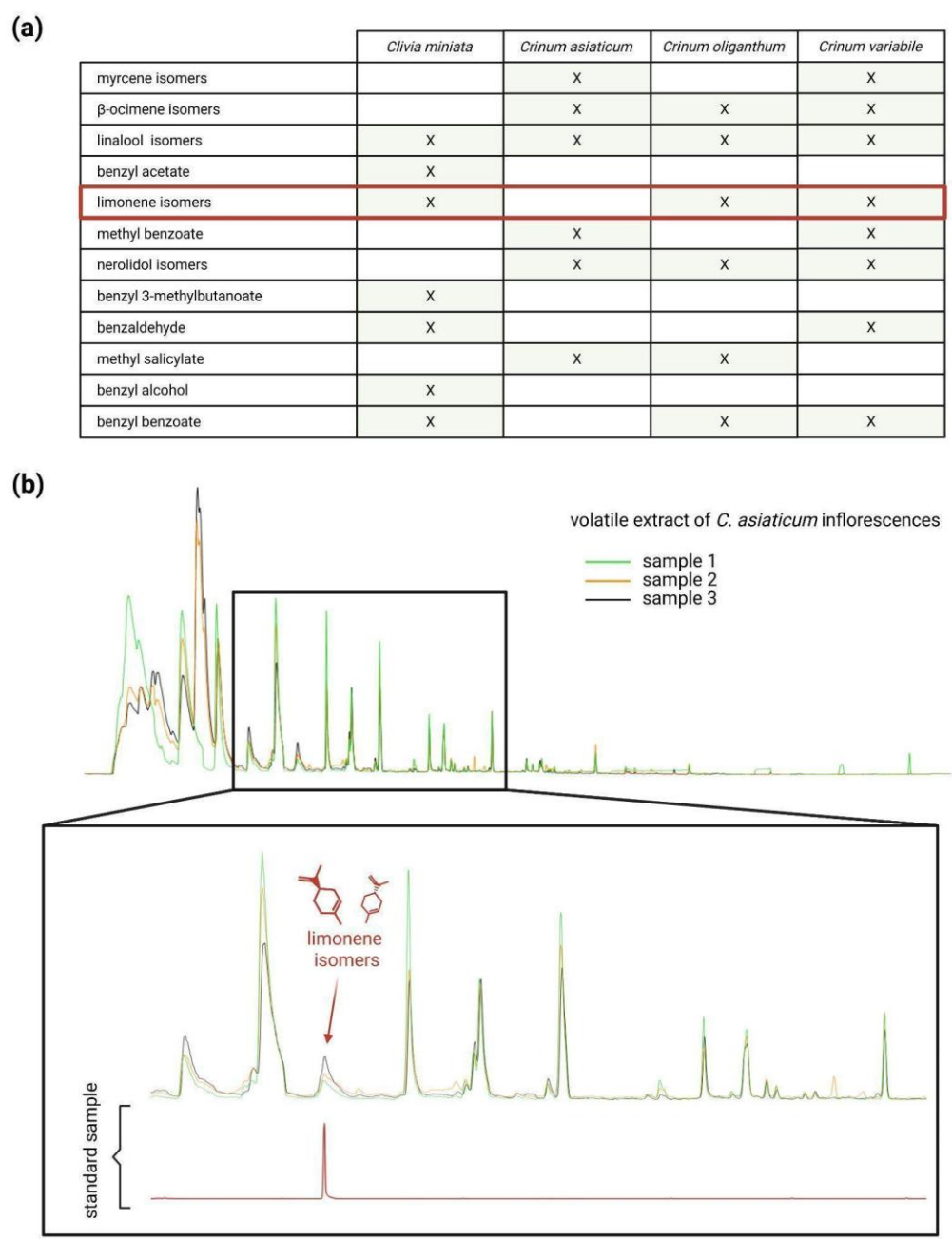

**Supplementary Figure 1. Limonene enantiomers are potential semiochemical cues guiding host plant location by adult *S. picta*.** (a) Reported floral volatiles from *Crinum* species and *C. miniata*, compiled from literature sources (Miyake et al. 1998; Manning and Snijman 2002; Báez et al. 2011, Kiepiel and Johnson 2014). Limonene enantiomers are highlighted in red. (b) Three GC traces of *Crinum asiaticum* inflorescence headspace are aligned with that of the synthetic limonene enantiomers (in red), as both enantiomers co-elute under the chromatographic conditions used. Sample 1 is shown in green, sample 2 in orange, and sample 3 in black.

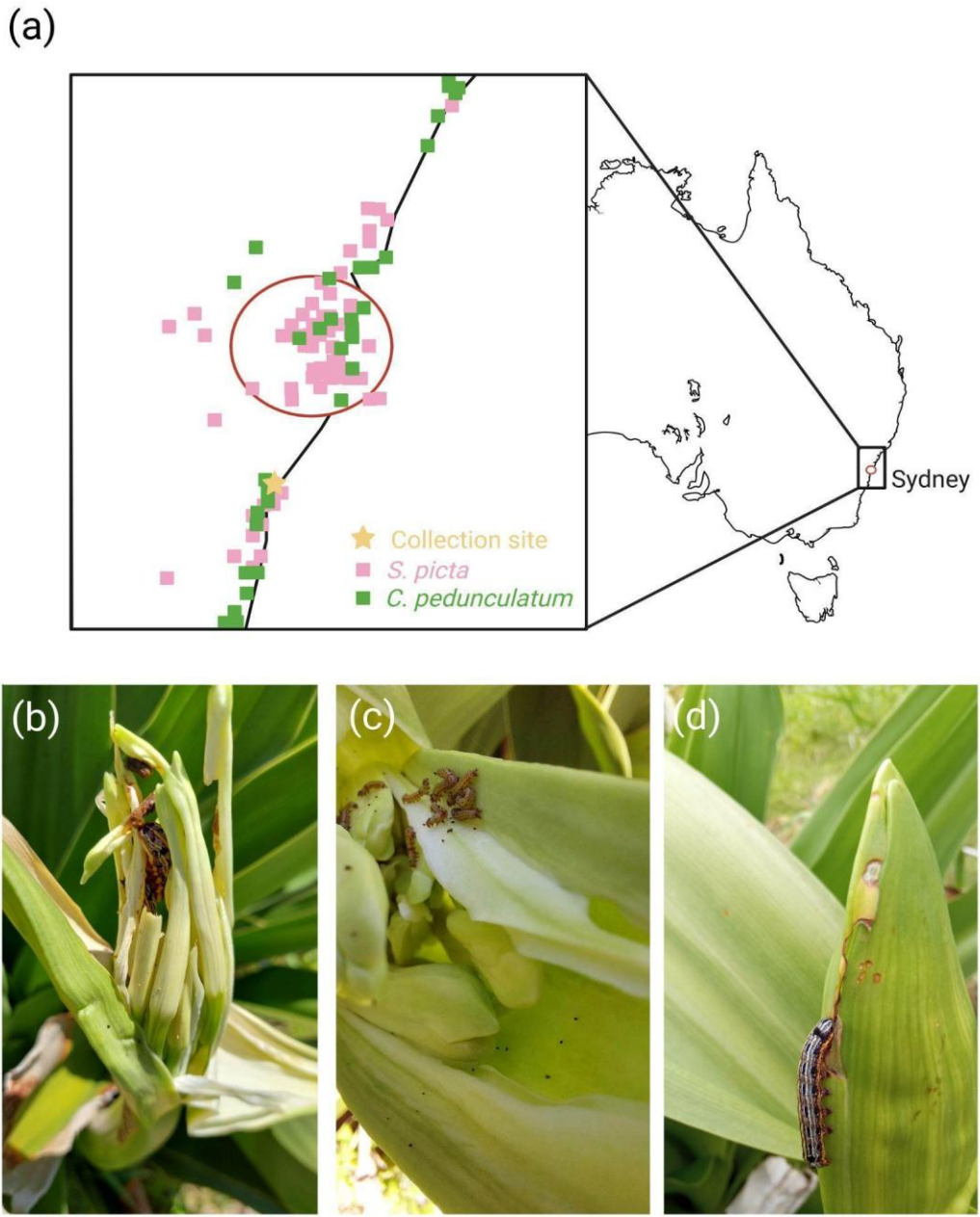

Supplementary Figure 2. Larvae of *S. picta* were collected from the inflorescence of *C. pedunculatum* in the Thirroul region, Australia. (a) Schematic representation of all recorded identifications of *S. picta* and *C. pedunculatum* made by iNaturalist users (Matheson CA 2014) between 2012 and 2022 around Sydney, Australia. Sites where *S. picta* was observed are marked in pink, while locations of *C. pedunculatum* are shown in green. The sample collection site (5 Hamilton Rd, Thirroul, NSW 2515, Australia) is indicated by a yellow star. (b-d) Photographs documenting the feeding behavior of *S. picta* at different larval stages. The images were taken at the collection site on *C. pedunculatum* (photo credit: A. Comte).

**Supplementary Table 2. 3D models generated by AlphaFold 2 exhibit high accuracy, particularly**
**in the binding regions of SpicORs, SlituORs, and SlitORs. The AlphaFold 3 web server did not**
**approve the accuracy of the models produced by the previous version.** The pLDDT scores
obtained for the full sequence and the binding regions of the 213 receptors studied are compiled for the
best models generated by the AlphaFold 2 and AlphaFold 3 web servers (Jumper et al. 2021; Abramson
et al. 2024). pLDDT scores of 90 or higher are colored dark blue, scores between 85 and 90 in light
blue, and scores between 76 and 85 in yellow.

|  | AlphaFold2 models accuracy (pLDDT score analysis) |  |  |  |  |  | AlphaFold3 models accuracy (pLDDT score analysis) |  |  |  |  |  |
| --- | --- | --- | --- | --- | --- | --- | --- | --- | --- | --- | --- | --- |
|  | *S. picta* |  | *S. litura* |  | *S. littoralis* |  | *S. picta* |  | *S. litura* |  | *S. littoralis* |  |
|  | Full sequence | Binding area | Full sequence | Binding area | Full sequence | Binding area | Full sequence | Binding area | Full sequence | Binding area | Full sequence | Binding area |
| OR1 | 85.692 | 87.442 | 85.262 | 88.180 | 86.054 | 88.018 | 82.475 | 87.978 | 81.956 | 87.898 | 86.858 | 88.166 |
| OR3 | 90.258 | 89.433 | 89.646 | 87.793 | 90.086 | 88.918 | 86.704 | 86.698 | 86.780 | 86.722 | 86.858 | 87.187 |
| OR4 | 89.462 | 91.683 | 89.840 | 92.239 | 90.943 | 91.655 | 85.731 | 88.880 | 86.047 | 88.383 | 86.171 | 87.449 |
| OR5 | 91.629 | 92.916 | 92.128 | 93.839 | 90.839 | 93.129 | 86.892 | 89.035 | 87.168 | 89.394 | 87.737 | 90.127 |
| OR6 | 90.048 | 92.120 | 90.668 | 92.341 | 90.030 | 91.094 | 85.555 | 86.200 | 84.695 | 87.143 | 85.405 | 86.043 |
| OR7 | 86.116 | 88.016 | 86.093 | 87.811 | 86.688 | 87.948 | 86.149 | 88.816 | 84.760 | 86.636 | 85.159 | 87.612 |
| OR8 | 88.096 | 91.169 | 88.098 | 90.475 | 88.227 | 88.865 | 84.374 | 86.239 | 83.279 | 85.921 | 84.047 | 85.195 |
| OR9 | 91.247 | 92.166 | 90.567 | 90.200 | 91.385 | 91.642 | 86.828 | 90.043 | 83.797 | 84.692 | 87.819 | 90.622 |
| OR10 | 89.549 | 91.051 | 89.767 | 90.971 | 89.636 | 90.753 | 84.981 | 87.553 | 84.764 | 86.727 | 83.891 | 86.677 |
| OR11 | 90.382 | 93.411 | 90.545 | 93.196 | 91.095 | 92.540 | 81.436 | 84.877 | 81.841 | 83.866 | 82.895 | 83.921 |
| OR12 | 91.584 | 93.862 | 91.435 | 93.281 | 91.145 | 93.479 | 85.886 | 90.218 | 87.123 | 90.763 | 86.985 | 90.406 |
| OR13 | 91.123 | 93.510 | 91.264 | 93.731 | 90.949 | 94.151 | 86.432 | 89.189 | 86.538 | 89.542 | 86.488 | 89.849 |
| OR14 | 87.500 | 87.896 | 87.708 | 87.783 | 88.281 | 87.917 | 81.138 | 83.401 | 81.893 | 83.101 | 81.697 | 83.494 |
| OR15 | 90.843 | 90.822 | 91.900 | 92.713 | 91.288 | 92.521 | 85.771 | 86.986 | 86.498 | 87.710 | 85.767 | 87.858 |
| OR16 | 91.636 | 94.639 | 91.588 | 93.663 | 91.565 | 93.837 | 85.070 | 85.679 | 84.582 | 85.316 | 84.985 | 85.463 |
| OR17 | 91.378 | 92.518 | 92.920 | 94.510 | 93.018 | 94.498 | 87.954 | 87.771 | 86.900 | 87.573 | 87.609 | 87.520 |
| OR18 | 91.398 | 89.734 | 91.697 | 90.561 | 91.215 | 90.279 | 85.664 | 84.083 | 85.342 | 84.062 | 85.173 | 83.226 |
| OR19 | 93.044 | 95.309 | 92.603 | 93.495 | 91.981 | 94.451 | 87.488 | 87.723 | 87.101 | 87.460 | 87.534 | 87.620 |
| OR20 | 89.591 | 92.434 | 89.416 | 92.114 | 89.740 | 90.451 | 84.796 | 85.576 | 85.378 | 86.763 | 84.392 | 84.431 |
| OR21 | 93.119 | 93.763 | 90.674 | 91.391 | 92.575 | 91.772 | 88.054 | 86.248 | 85.808 | 82.826 | 87.259 | 85.364 |
| OR22 | 91.578 | 91.820 | 91.434 | 91.642 | 91.465 | 92.306 | 86.390 | 85.860 | 87.177 | 86.705 | 87.190 | 86.294 |
| OR23 | 89.526 | 93.151 | 90.117 | 94.212 | 89.935 | 94.422 | 86.506 | 90.568 | 86.653 | 91.165 | 86.260 | 90.844 |
| OR24 | 89.496 | 92.318 | 89.995 | 92.305 | 89.217 | 91.151 | 86.366 | 87.356 | 86.646 | 87.743 | 86.384 | 87.611 |
| OR25 | 89.288 | 91.569 | 88.572 | 90.415 | 88.386 | 90.365 | 86.408 | 86.287 | 86.188 | 86.374 | 86.378 | 86.190 |
| OR26 | 93.797 | 96.014 | 93.007 | 95.626 | 92.434 | 95.868 | 87.927 | 89.752 | 87.197 | 88.423 | 87.469 | 89.159 |
| OR27 | 86.639 | 89.181 | 86.030 | 86.823 | 86.617 | 89.161 | 82.629 | 84.921 | 82.890 | 84.612 | 82.947 | 84.609 |
| OR28 | 88.685 | 91.587 | 87.891 | 91.466 | 87.122 | 89.150 | 79.863 | 85.708 | 78.596 | 85.778 | 79.637 | 86.807 |
| OR29 | 90.605 | 91.010 | 90.874 | 91.223 | 90.850 | 91.366 | 83.988 | 85.070 | 85.044 | 85.599 | 84.353 | 85.413 |
| OR30 |  |  | 91.321 | 91.466 | 92.004 | 92.692 |  |  | 84.775 | 85.596 | 85.893 | 87.619 |
| OR31 | 88.321 | 90.360 | 89.009 | 91.520 | 88.701 | 90.465 | 81.555 | 84.629 | 83.186 | 85.243 | 83.076 | 84.931 |
| OR32 | 89.550 | 89.284 | 89.828 | 89.582 | 89.080 | 88.867 | 87.281 | 87.482 | 86.887 | 87.401 | 86.190 | 86.281 |
| OR33 | 89.539 | 90.297 | 90.139 | 90.832 | 90.533 | 91.082 | 86.115 | 84.767 | 86.374 | 84.551 | 86.346 | 85.045 |
| OR34 | 88.778 | 90.308 | 88.336 | 90.240 | 87.419 | 88.604 | 85.818 | 85.764 | 85.899 | 85.096 | 86.659 | 86.317 |
| OR35 | 91.712 | 91.800 | 91.186 | 92.093 | 91.147 | 92.387 | 85.724 | 88.640 | 85.321 | 88.142 | 85.220 | 88.380 |
| OR36 | 92.861 | 93.522 | 92.973 | 93.357 | 92.933 | 93.496 | 84.104 | 87.830 | 84.554 | 87.907 | 84.359 | 87.712 |
| OR37 | 91.993 | 92.480 | 92.252 | 93.792 | 92.018 | 93.037 | 88.233 | 85.599 | 88.494 | 85.543 | 88.541 | 85.031 |
| OR38 |  |  | 91.197 | 91.610 | 91.502 | 92.363 |  |  | 87.682 | 89.682 | 87.325 | 89.486 |
| OR39 | 90.763 | 93.452 | 91.507 | 93.705 | 87.190 | 88.396 | 86.958 | 90.928 | 86.347 | 89.779 | 83.702 | 84.243 |
| OR40 | 88.649 | 88.761 | 88.556 | 89.429 | 88.522 | 89.076 | 83.820 | 82.762 | 83.454 | 82.233 | 83.769 | 83.243 |
| OR41 | 93.118 | 92.599 | 92.620 | 91.601 | 93.113 | 93.156 | 84.323 | 85.266 | 85.604 | 85.443 | 84.787 | 85.414 |
| OR42 | 87.916 | 90.696 | 87.121 | 89.502 | 87.345 | 90.072 | 84.255 | 88.229 | 83.710 | 86.333 | 84.076 | 87.876 |
| OR43 | 89.252 | 86.989 | 89.752 | 88.320 | 88.715 | 88.615 | 85.341 | 84.000 | 85.064 | 84.873 | 83.273 | 85.301 |
| OR44 | 88.413 | 88.723 |  |  | 88.753 | 88.523 | 84.263 | 83.213 |  |  | 84.868 | 83.084 |
| OR45 | 87.551 | 87.622 | 87.684 | 83.468 | 87.489 | 88.267 | 84.106 | 83.345 | 82.853 | 79.016 | 82.944 | 81.609 |
| OR46 | 90.253 | 89.549 | 89.731 | 85.700 | 89.534 | 86.710 | 84.957 | 85.093 | 84.615 | 82.123 | 84.920 | 83.704 |
| OR47 | 91.328 | 93.315 | 91.810 | 94.136 | 91.840 | 93.690 | 84.184 | 89.338 | 85.380 | 90.457 | 85.516 | 90.339 |
| OR48 | 91.276 | 89.551 | 91.549 | 90.217 | 91.515 | 89.369 | 84.672 | 84.029 | 84.861 | 85.411 | 84.863 | 84.852 |
| OR49 | 91.411 | 89.612 | 90.656 | 88.877 | 91.335 | 89.144 | 85.161 | 85.377 | 85.448 | 85.525 | 85.240 | 84.442 |
| OR50 | 90.678 | 88.199 | 91.291 | 88.827 | 90.664 | 89.466 | 85.887 | 82.392 | 86.423 | 84.752 | 87.075 | 86.682 |
| OR51 | 91.288 | 92.054 |  |  | 91.601 | 92.580 | 86.442 | 86.955 |  |  | 86.193 | 86.823 |
| OR52 | 92.681 | 94.437 | 92.229 | 94.367 | 91.922 | 94.188 | 86.745 | 90.318 | 86.644 | 90.258 | 86.005 | 89.364 |
| OR53 | 89.972 | 90.356 | 90.114 | 91.188 | 90.129 | 90.697 | 85.529 | 87.200 | 85.202 | 86.775 | 85.576 | 86.519 |
| OR54 | 89.464 | 90.341 | 88.902 | 90.293 | 91.135 | 92.012 | 83.914 | 86.275 | 84.061 | 85.551 | 84.325 | 86.251 |
| OR55 | 92.659 | 93.545 | 92.558 | 93.579 | 92.729 | 93.800 | 85.213 | 88.333 | 85.200 | 89.171 | 85.635 | 89.436 |
| OR56 | 90.689 | 93.831 | 90.552 | 92.915 | 90.313 | 93.648 | 83.818 | 84.950 | 84.152 | 84.612 | 84.093 | 85.175 |
| OR57 | 91.049 | 92.181 | 90.012 | 93.821 | 89.262 | 93.144 | 85.069 | 86.538 | 84.799 | 86.915 | 85.583 | 87.319 |
| OR58 | 82.552 | 89.062 | 82.021 | 89.582 | 82.124 | 87.450 | 77.558 | 85.343 | 76.891 | 83.634 | 77.763 | 86.331 |
| OR59 | 90.030 | 91.266 | 90.155 | 91.282 | 89.474 | 90.540 | 85.527 | 85.553 | 85.707 | 86.738 | 85.475 | 86.063 |
| OR60 | 88.304 | 90.738 | 88.723 | 89.599 | 89.432 | 90.131 | 80.812 | 81.183 | 81.815 | 82.827 | 81.754 | 81.909 |
| OR61 | 90.447 | 92.446 | 90.434 | 91.455 | 90.746 | 92.738 | 82.237 | 83.856 | 82.532 | 84.887 | 82.288 | 85.092 |
| OR63 | 88.116 | 89.456 | 88.578 | 89.596 | 87.631 | 87.314 | 79.271 | 82.557 | 80.010 | 81.863 | 79.127 | 79.833 |
| OR64 | 90.765 | 90.289 |  |  | 90.552 | 91.707 | 87.024 | 87.437 |  |  | 87.221 | 87.414 |
| OR65 | 86.181 | 85.062 | 86.556 | 90.026 | 86.244 | 87.271 | 80.864 | 88.523 | 80.082 | 86.908 | 81.720 | 88.013 |
| OR66 | 92.747 | 94.227 | 92.628 | 94.009 | 92.732 | 93.898 | 87.164 | 88.237 | 87.586 | 88.529 | 87.204 | 88.725 |
| OR67 | 89.099 | 90.829 | 88.999 | 91.583 | 89.326 | 91.968 | 83.525 | 86.295 | 84.109 | 87.388 | 83.722 | 87.534 |
| OR68 | 88.506 | 91.338 | 87.548 | 89.957 | 87.045 | 90.048 | 83.026 | 87.868 | 82.972 | 87.727 | 82.242 | 86.257 |
| OR69 | 88.649 | 90.707 | 88.042 | 89.796 | 88.876 | 90.916 | 85.991 | 88.350 | 86.019 | 87.270 | 86.124 | 87.382 |
| OR70 | 85.595 | 89.986 | 86.432 | 89.416 | 85.726 | 89.910 | 82.151 | 88.463 | 82.132 | 87.100 | 81.939 | 87.512 |
| OR71 | 91.580 | 93.328 | 92.114 | 93.616 | 92.069 | 94.318 | 84.831 | 86.206 | 85.177 | 86.721 | 85.081 | 86.637 |
| OR72 |  |  | 91.782 | 95.314 | 92.306 | 94.263 |  |  | 85.442 | 88.617 | 85.064 | 87.039 |
| OR73 | 92.441 | 92.968 | 93.272 | 93.323 | 91.764 | 92.633 | 87.344 | 88.855 | 87.201 | 88.950 | 86.802 | 88.898 |
| OR74 | 86.461 | 88.387 | 86.271 | 89.128 | 86.349 | 88.033 | 83.174 | 86.821 | 82.925 | 86.797 | 83.190 | 86.529 |
| OR75 | 91.160 | 93.222 | 90.931 | 93.247 | 90.484 | 93.029 | 87.494 | 90.534 | 87.442 | 90.112 | 87.146 | 89.912 |
| OR76 |  |  | 87.875 | 90.488 |  |  |  |  |  | 84.128 | 86.931 |  |
|  |  |  |  |  |  |  |  |  |  |  | </ |  |

pLDDT score ≥ 90 90 > pLDDT score ≥ 85 85 > pLDDT score ≥ 76

### **References**

- Abramson J, Adler J, Dunger J, Evans R, Green T, Pritzel A, Olaf Ronneberger O, Willmore L, Ballard
  AJ, Bambrick J et al. 2024. Accurate structure prediction of biomolecular interactions with AlphaFold 3.
  Nature 630(8016):493-500. [10.1038/s41586-024-07487-w](https://doi.org/10.1038/s41586-024-07487-w).
53 Báez D, Pino JA, Morales D. 2011. Scent Composition in Some Cuban Flowers: *Clitoria Fairchildiana*
  RA Howard, *Brunfelsia Nitida* Benth. and *Crinum Oliganthum* Urban. Journal of Essential Oil Bearing
  Plants 14(4):383-386. [10.1080/0972060X.2011.10643590](https://doi.org/10.1080/0972060X.2011.10643590).
56 Bouysset C, Fiorucci S. 2021. ProLIF: a library to encode molecular interactions as fingerprints. Journal
  of cheminformatics 13(1):72. [10.1186/s13321-021-00548-6](https://doi.org/10.1186/s13321-021-00548-6).
58 Comte A, Lalis M, Brajon L, Moracci R, Montagné N, Topin J, Jacquin-Joly E, Fiorucci S. 2025.
  Accelerating ligand discovery for insect odorant receptors. International Journal of Biological Sciences
  21(5):2101. [10.7150/ijbs.105648](https://doi.org/10.7150/ijbs.105648).
61 DeLano WL. 2002. Pymol: An open-source molecular graphics tool. CCP4 Newsl. Protein Crystallogr
  40(1):82-92.
- Jumper J, Evans R, Pritzel A, Green T, Figurnov M, Ronneberger O, Tunyasuvunakool K, Bates R,
  Žídek A, Potapenko A et al. 2021. Highly accurate protein structure prediction with AlphaFold. Nature
  596(7873):583-589. [10.1038/s41586-021-03819-2](https://doi.org/10.1038/s41586-021-03819-2).
66 Kiepiel I, Johnson SD. 2014. Shift from bird to butterfly pollination in *Clivia* (Amaryllidaceae). *American*
  *Journal of Botany* 101(1):190-200. [10.3732/ajb.1300363](https://doi.org/10.3732/ajb.1300363).
68 Manning JC, Snijman D. 2002. Hawkmoth-pollination in *Crinum variable* (Amaryllidaceae) and the
  biogeography of sphingophily in southern African Amaryllidaceae. South African Journal of Botany
  68(2):212-216. [doi.org/10.1016/S0254-6299(15)30422-1](https://doi.org/10.1016/S0254-6299%2815%2930422-1).
71 Matheson CA. 2014. inaturalist. *Reference Reviews* 28(8):36-38. [10.1108/RR-07-2014-0203](https://doi.org/10.1108/RR-07-2014-0203).
72 Miyake T, Yamaoka R, Yahara T. 1998. Floral scents of hawkmoth-pollinated flowers in Japan. Journal
  of Plant Research 111(2):199-205. [10.1007/BF02512170](https://doi.org/10.1007/BF02512170).
74 Wang H, Song J, Hunt BJ, Zuo K, Zhou H, Hayward A, Li B, Xiao Y, Geng X, Bass C, et al. 2024a.
  UDP-glycosyltransferases act as key determinants of host plant range in generalist and specialist
  *Spodoptera* species. Proceedings of the National Academy of Sciences 121(19), e2402045121.
  [10.1073/pnas.2402045121](https://doi.org/10.1073/pnas.2402045121).
